# Isolation and introductions disrupt the homogeneity of Argentine ants in Europe

**DOI:** 10.64898/2026.06.09.731050

**Authors:** Srikrishna Narasimhan, Enikő Csata, Laura Trapero-Camesella, Marion Cordonnier, Giuseppe Mannino, Luca Pietro Casacci, Magdalena Witek, Iago Sanmartín-Villar

## Abstract

The introduction of alien species into new habitats stands as a pressing economic and ecological challenge but it is also essential for unveiling evolutionary processes. The introduction of the Argentine ant (*Linepithema humile*) led to the spread of a single supercolony through different continents and thousands of kilometres like in Europe, from Northwest Spain to Greece. It was assumed that the high invasiveness of the species mainly relied on the lack of agonism among colonies, an effect derived from its introduction. However, recent studies suggest that local adaptations and evolutionary divergence could involve the disruption of the Argentine ant “empire” into a mosaic of competitive colonies. We investigated how isolation affects population divergence by comparing mainland and island populations in two distant regions colonized in Spain and in Greece with morphology, agonism, cuticular hydrocarbons, and genetic diversity of ant workers. Our results showed that all colonies sampled belonged to the most spread supercolony in Europe (main supercolony) except one sampled in Crete (Heraklion; Greece), which resulted to be a supercolony not registered in Europe. The Heraklian supercolony showed a different chemical and genetic profile and hostile agonism towards the other Greek colonies. Differences between islands and mainland colonies belonging to the main supercolony were higher in Galiza than in Greece. Surprisingly, the chemical profile of the Cretan colony belonging to the main supercolony showed more similarity with the Galizan colonies than with the Greek mainland, suggesting that *L. humile* may have been introduced into Greece through this island instead of the mainland. Our study suggests that local adaptations in Argentine ant colonies can trigger competition between colonies. Our data strongly support the existence of a candidate supercolony which highlights either ongoing introductions of *L. humile* in Europe or gaps in our understanding of its metapopulation dynamics.

## Introduction

Species dispersal out of their native distribution constitutes one of the main ecological impacts nowadays (Sherpa and Després, 2021; Sesay *et al*., 2024). Species are intentionally and unintentionally transported due to the worldwide resource exchange facilitated by globalisation (Hulme *et al*., 2008) or they spread out of their distributions following the environmental conditions caused by current climate change (e.g. Urban, 2020). Some of the introduced species adapt, reproduce and spread through the new areas negatively affecting the ecological relationships and the environment, *i.e.* becoming invasive (Pyšek *et al*., 2020). Despite the ecological damage caused by biological invasions, this process also provides an excellent opportunity to observe rapid evolutionary changes. Most species struggle or fail to survive under changing environmental conditions (Sorte *et al*., 2013), whereas many invasive species succeed in overcoming these challenges and come to dominate ecological interactions in the areas they were introduced. Among exotic invasive species, insects show several biological traits facilitating invasiveness (Zhao *et al*., 2023). However, some social species, such as ants, possess an additional trait that boosts their competence against native species: formation of supercolonies. Understood as the lack of aggressiveness among colonies of unicolonial species (Tsutsui *et al*., 2000; Heikki Helanterä, 2022), *i.e.* those organised in interconnected nests, the existence of supercolony reduces intraspecific competition and thus, increases the ecological dominance for resources, frequently causing the displacement of native species (Eyer *et al*., 2021).

The worldwide adaptation (Wetterer *et al*., 2009; Angulo *et al*., 2024) of the argentine ant (*Linepithema humile* Mayr 1868) is assumed by its particular aggression pattern and its supercolonial behaviour. Argentine ants show high interspecific aggression in its native distribution (Holway and Suarez, 2004; LeBrun *et al*., 2007) and when introduced in a new region (*e.g.* Touyama, et al., 2003; Trigos-Peral, et al., 2021; Sanmartín-Villar, et al., 2022). However, the agonistic intraspecific interactions that are considered as one of the factors limiting the distribution of short-range supercolonies in its native area (Heller, 2004), seem to disappear when introduced by processes as founder and bottleneck effects (Tsutsui *et al*., 2000; Suarez, Holway and Case, 2001), producing long-range supercolonies (Giraud 2002; but see Seko et al., 2021). Different long-range *L. humile* supercolonies were discovered in their introduced areas: at least seven supercolonies were identified in California (Thomas 2005, 2006, 2007), multiple supercolonies in Florida (Buczkowski et al., 2004), two in South Africa (Mothapo and Wossler, 2011), and four in Japan (Sunamura *et al*., 2009; Hayasaka *et al*., 2023). Three supercolonies were identified in Europe, where *L. humile* was detected in almost every country located in the southern part of the continent: the main supercolony, present also in other continents (Suhr et al., 2011; Van Wilgenburg et al., 2010) spread from the coast of NW Spain to Greece (Gómez & Espadaler 2005); the Catalunyan supercolony, limited to the Spanish Mediterranean coast (Giraud 2002) and Balearic islands (Sara Castro-Cobo *et al*., 2021); and the Corsican supercolony (Blight *et al*., 2012), limited to the French Mediterranean coast and Corsica. Differences among supercolony traits and the strong agonistic response when individuals from different supercolonies encounter each other suggests that several introduction events had occurred in these areas.

In opposition to the pattern observed previously, certain colonies that belong to the main supercolony showed genetic variability and agonistic interactions in Galiza (NW Spain), even surpassing the values observed between the main and the Corsican supercolonies (Sanmartín-Villar *et al.,* 2022). This phenomenon suggests that the presumed vast “empire” of some supercolonies is not as homogeneous as assumed, and that this internal variability had not been detected in previous studies (Tsutsui *et al*., 2000; Suarez, 2001; Giraud, 2002). In that case, the supercolony could be composed of a mosaic of relatively tolerant colonies in which the outcome of their interactions depends on the genetic or recognition cue similarities or the individual behavioural variability. On the other hand, the variability within the same supercolony could be explained as a process of evolutionary divergence. The local adaptation and isolation over the two decades between studies performed by Giraud et al. and Sanmartín-Villar *et al*. (2000–2020) may have contributed to the re-emergence of inter-colony aggression that had been reduced during introduction. The mechanism behind this process seems key to understanding short evolutionary processes that might be essential in a changing environment.

Our main goal was to unravel the potential divergence in allopatric populations based on the dispersal dynamics of *L. humile*. We considered two levels of isolation: due to local geographical barriers and distant geographical locations. To analyse isolation due to local geographical barriers, we analysed populations from the coastal mainland and maritime islands. The interest in performing this analysis relies on the intraspecific diversity observed in areas related to island systems in South Europe (Giraud, 2002; Blight *et al*., 2010; Sarnat and Moreau, 2011; Sanmartín-Villar, Silva, *et al*., 2022). Although *L. humile* occurs on multiple islands, active dispersal between island and mainland is likely limited, as only males disperse by flight (Passera and Keller, 1994). To analyse isolation patterns over distant geographical locations, we compared populations from the longitudinal extremes of the European distribution: South West (Galiza, NW Spain) and South East Europe (Greece). Although individuals from NW Spain and North Italy (Giraud et al., 2002) or Greece (Gómez & Espadaler, 2005) belong to the same supercolony and showed no aggression between them, we considered that individuals of colonies separated over long distances were not connected because it would involve a travel of thousands of kilometres. *Linepithema humile* was most likely passively dispersed throughout Europe with the earliest European records in Portugal (Wetterer *et al*., 2009). In this context, the comparison of the selected sampled areas could determine the effect of isolation in shaping colony dynamics through distant populations of an assumed single supercolony.

We performed a four-axis study focused on morphometrics, agonistic behaviour, cuticular hydrocarbon profile and genetic variability (parameters previously studied in the divergence of ant supercolonies, (Casadei-Ferreira, Feitosa and Pie, 2022) and Van Wilgenburg *et al*., 2010) to obtain a holistic view of the effect on population dynamics due to isolation on *Linepithema humile* main supercolony. We predicted differences in the four axes between islands and mainland populations and between Galiza and Greece due to allopatry, local adaptations and divergence.

## Materials and Methods

### Ant sampling and maintenance

A total of eight islands and seven mainland populations were sampled between October 2023 (Greece) and March 2024 (Galiza; Table 1). Ants from the Catalunyan supercolony (Sant Cugat; Table 1) were collected on 30 July 2024 and 5 April 2025 to compare their genetic and chemical variability with previously collected samples. Sampled localities were selected based on previous records (Galiza: Fraga-Cimadevila et al., 2025; Sanmartín-Villar et al., 2022; Greece: Salata et al., 2019; Stephen Clifford (https://www.inaturalist.org/observations/181209537); Catalunya: Xavier Espadaler, pers. comm. 2021) or according to the relevance of their geolocation and the goals of the study. Galizan samples were collected again on 29 August 2025 to control for the seasonal effect of agonism (Sanmartín-Villar, et al., 2022; Supplementary Material).

**Table 1.** Populations sampled. Rows are ordered following a longitudinal and latitudinal gradient. *Dist*: shorter distance, in km, between the mainland and each island (measured with Google Earth). *Location*: place where the ants were found. *Day*: day on which ants were sampled. *Maintenance*: number of days of maintenance in captivity from ant sampling to the agonism test. *Ref*: previous records followed for sampling (1: Narasimhan et al., 2026; 2: Fraga-Cimadevila et al., 2025; 3: Sanmartín-Villar, 2022; 4: Xavier Espadaler, personal communication (2021); 5: Stephen Clifford through *iNaturalist* (https://www.inaturalist.org/observations/181209537; 2023); 6: Salata et al., 2019. *: Arousa and Toxa islands are connected to the mainland by bridges. ^§^: samples used for CHC determination. ^§§^: samples used for microsatellite determination. No behavioral tests were performed with Catalunyan supercolonies, so ants were not kept in the laboratory.

| Area | Locality | Code | Type | Dist | Habita<br>t | Location | Latitude | Longitude | Altitude<br>(m) | Base<br>d on | Day | Mainte<br>nance |
| --- | --- | --- | --- | --- | --- | --- | --- | --- | --- | --- | --- | --- |
| Greec<br>e | Loutraki | LOU | Mainland | - | Garde<br>n | Pine base | 37°57'48"<br>N | 22°58'19"E | 4 | 3 | 28/10/2023 | 4 |
| Greec<br>e | Athens | ATH | Mainland | - | Park | Trees,<br>rocks | 37°58'27"<br>N | 23°44'08"E | 93 | 4 | 27/10/2023 | 5 |
| Greec<br>e | Heraklion<br>(Crete) | HRK | Island | 95.86 | Garde<br>n | Tiles | 35°20'14"<br>N | 25°07'50"E | 23 | - | 21/10/2023 | 11 |
| Greec<br>e | Hersoniso<br>s (Crete) | HSS | Island | 95.86 | Beach | Carton<br>board | 35°20'04"<br>N | 25°23'02"E | 11 | 3 | 20/10/2023 | 12 |
| Galiza | Cortegada | COR | Island | 0.19 | Forest<br>beach | Pine base | 42°36'60"<br>N | 8°46'58"W | 9 | 1 | 10-12/03/2024 | 12-14 |
| Galiza | Carril | CAR | Mainland | - | Park | Residuals | 42°37'15"<br>N | 8°46'23"W | 17 | 2 | 10-12/03/2024 | 12-14 |
| Galiza | Arousa | ARO | Island | 1.69* | Garde<br>n | Compost | 42°33'43"<br>N | 8°52'36"W | 9 | 1 | 10-12/03/2024 | 12-14 |
| Galiza | Vilanova | VIL | Mainland | - | Park | Rock | 42°34'01" N | 8°49'43"W | 30 | 1&2 | 10-12/03/2024 | 12-14 |
| Galiza | Toxa | TOX | Island | 0.32* | Park | Pine base | 42°29'12" N | 8°50'55"W | 33 | 1 | 10-12/03/2024 | 12-14 |
| Galiza | Lanzada | LAN | Mainland | - | **Coast** | **Stones** | **42°25'49" N** | **8°52'22"W** | **5** | - | 10-12/03/2024 | 12-14 |
| Galiza | Refoxos | REF | mainland | - | Park | Residuals | 42°35'45.09"N | 8°44'38.79" W | 37 | - | 01/05/2023 | 0 |
| **Catalunya** | **Sant Cugat** | **SCU** | **Mainland** | - | **Park** | **Sandy soil** | **41°28'28" N** | **2°04'38"E** | **147** | 5 | 30/07/2024<sup>§</sup><br>§ &<br>05/04/2025<sup>§</sup> | 0 |

*Linepithema humile* nests were searched by digging the soil with garden shovels and by picking rocks, stones, tiles, logs, and garbage by two (Galiza) or three (Greece) observers. Six nests were found in Galiza (three mainland and three island populations) and four nests from Greece (two mainland and two island populations). Ant workers were collected using brushes and pooters. Around 50 and 24 workers were preserved in ethanol (96%) from each nest for morphometric and genetic analyses respectively. Around 2000 workers were transferred along with nest soil to plastic containers whose interiors were firstly coated with fluon (polytetrafluoroethylene; ITAflon©) to avoid the escape of ants for agonism tests (240 and 150 ants from each nest from Galiza and Greece respectively). Wet cotton and *ad libitum* honey were provided to each container to keep the ants alive until the agonism test were performed (Table 1). Around 20 ants of each colony were frozen at -80°C (Greece: 07/11/2023; Galiza: 18/04/2024) for the analysis of the cuticular hydrocarbons.

### Morphometrics

To analyse if local adaptation and isolation triggers morphological differences, the heads of 50 individuals of each colony (N_Greece_= 200; N_Galiza_= 300) were analysed under the binocular lens (Olympus SZ60; Olympus SZH10). We measured the eye span, the maximum head width, and the distance between the clipeus base and the occipucium as face length as a proxy of body size (Gibb and Parr, 2013). Statistical analyses were performed using R (www.r-project.org). A one-way multivariate analysis of variance (MANOVA) was conducted to assess the effect of population on three morphological traits (eyespam size, width, and height). Following a significant MANOVA, separate univariate ANOVAs were performed for each trait, followed by Tukey’s post-hoc test (TukeyHSD; Miller, 1981) for pairwise comparisons.

### Agonism test

To examine individual recognition of ants, we conducted one-on-one confrontations between workers from all possible colony pairs coming from the same geographical region (N_Greece_= 6; N_Galiza_= 15). Tests were performed within an average of 10 days in Greece and 11 days in Galiza from the date of collection (Table 1). We did not confront Greek and Galizan ant workers due to the temporal distance of their sampling and possible subsequent behavioural differences resulting from captivity (Dittmann et al., 2019; but see Giraud *et al*. 2002). Control confrontations were composed of two ant workers from the same colony (N_Greece_= 4; N_Galiza_= 6). Each confrontation was replicated 30 times with new ants from the same colony over two consecutive days between 10:00h and 14:00h (N_Greece_= 300 dishes containing 600 ant workers; N_Galiza_= 630 dishes containing 1260 ant workers).

Ant workers were individually transferred from their maintenance container to fluon-coated Petri dishes (Ø= 5.7 cm) to observe one-on-one confrontations. To avoid the resident effect (Vilela and Howse, 1986), the observers (Greece: 3; Galiza: 2) placed the ants in an alternative order in every next dish. Experimental individuals were randomly selected from a mix of individuals found in different parts of each experimental colony to avoid biasing their selection according to their age and/or task (potentially immature/nurses inside the nest and mature/foragers outside the soil). All selected workers possessed all their appendages. To record behavior from the initial interaction and standardize observation time to 10 minutes, ants were first recorded from the moment the first ant was added to the first dish. Recording continued until 10 minutes after the second ant was added to the final dish. The recording was done with three cameras (two Sony FDR-AX53 and Canon Legria HF M56) placed vertically and perpendicular to the dish. Each camera recorded 15 dishes at a time. The duration and the number of agonistic interactions among paired ants of each dish were analysed by one observer. Interactions were considered agonistic if they caused a change in direction of the ant during head-on interaction, pushed an ant away from them, bit any of the legs or antennae, flexed the gaster or bit each other with gaster flexion.

Assumptions for Normality were tested in both variables. In case of non-Normality, non-parametric analyses were performed. The effect of the kind of combination (control, mainland-mainland populations, island-island populations, island-mainland populations) and the region (Galiza, Greece) over the number and duration of the agonistic interactions were analysed as fixed factors in general linear mixed models considering Gamma distribution (*glmmTMB*, Brooks et al., 2017). The number of the Petri dish in which ants were analysed was added as a random factor. The number and duration of agonistic interactions were analysed among all possible combinations performed by Kruskal-Wallis tests. Means and standard errors are shown as x̄±SE.

### Cuticular hydrocarbon profile

To analyse the effect of allopatry on the ant cuticular hydrocarbon profile, three replicates of three ants from the same colony (9 ants from each colony) were extracted using 200 μL of hexane for 60 seconds. The extract was then concentrated under a gentle stream of nitrogen and dried, followed by re-suspension in 20 μL of hexane. Subsequently, 5 μL of the extract was injected into an on-column injector of an Agilent Technologies 6890 gas chromatograph, equipped with a single quadrupole mass spectrometer for analysis. The chromatographic separation was performed using a HP-5 capillary column (25 m × 0.2 mm × 0.33 μm film thickness). The temperature program was set as follows: initially held at 100°C for 5 min, followed by a linear increase of 10°C per minute up to 280°C, which was maintained for 50 min. During the analysis, the detector temperature was set at 300°C, the injector temperature at 280°C, and the helium carrier gas flow rate was 0.28 m/s. Peak areas and concentrations were determined using an electronic integrator. Mass spectrometry was conducted at 70 eV, with the injector and ion source temperatures set at 230°C and 250°C, respectively. Cuticular hydrocarbons were identified with the help of injections of standard series of odd n-alkanes, by calculating the equivalent chain length and by inspecting the mass spectrum fragmentation patterns. The areas under the peaks in the chromatogram were used for quantification, employing an external calibration curve constructed with n-tricosane as a reference compound.

### Cuticular hydrocarbon analysis

Chemical profiles were analysed as a matrix of absolute CHC peak abundances after log(x+1) transformation to reduce the influence of the largest peaks. Euclidean distances were then computed and used consistently across ordination and hypothesis testing. Differences among populations were first visualised by non-metric multidimensional scaling (NMDS) implemented with the function metaMDS (*vegan;* Oksanen et al., 2025), using two dimensions and 100 iterations. The quality of the ordination was evaluated from the stress values returned by the model.

To test for compositional differences we applied PERMANOVA (*adonis2*, *vegan;* Oksanen et al., 2026) with 9,999 permutations, considering as factors the locality of collection, the broader geographical areas (Galiza, Catalunya, Greece), and clusters identified in the NMDS ordination. Post-hoc pairwise contrasts were conducted with *pairwise.adonis2* (*pairwiseAdonis;* Arbizu, 2020). Because only three replicates were available per locality, tests at the locality level had limited statistical power and rarely yielded significant outcomes.

Finally we compared the distributions of Euclidean distances among selected groups of samples using two-sided Mann–Whitney U tests (*wilcox.test*, *stats;* Rey and Neuhäuser, 2011).

### Study of genetic structure of populations

To analyse the effect of allopatry on genetic differentiation we extracted DNA from 24 individuals from each colony (N_Greece_= 4; N_Galiza_= 6; N_Catalunya_= 1). DNA was extracted by homogenizing the thorax and legs of ants in a solution of 100 μl 5% chelex. The samples were firstly incubated at 56°C for 20 minutes, secondly at 95 °C for 10 min, and then centrifuged at 13000 rpm for 10 min. Then, 40μl of supernatant was stored at -20°C. Ants were assayed at 11 microsatellite markers: *Lhum-11*, *Lhum-13*, *Lhum-19*, *Lhum-28*, *Lhum-35*, *Lhum-39*, *Lhum-52*, *Lhum-62* (Butler *et al*., 2014), *LihuM1*, *LihuT1*, *LihuS3* (Ingram and Palumbi, 2002). Three sets of multiplex reactions were used with the forward primers labeled. The PCRs were performed in a total volume of 15 μl composed of 1.5 μl of DNA template, Multiplex PCR Master Mix (Qiagen), water and primers. For PCR amplification, a thermal cycler (Applied Biosystems) was used with the following PCR profile: 94°C for 15 min., then 35 cycles of: 95°C for 30 s, 57°C or 60°C (depending on the multiplex) for 30 s, 72 °C for 30s, with the final elongation of 72°C for 2min. PCR products were run on an ABI 3500 XL automated sequencer with the GeneScan™ 600 LIZ® Size Standard and analyzed using GENEMAPPER 4.1 (Applied Biosystems).

Three loci not polymorphic or suffering massive missing data (*Lhum_62*, *LihuM1*, *LihuS3*) were excluded from the analyses. The remaining eight loci were polymorphic and showed substantial variation (allele numbers: *Lhum-13*: 8; *Lhum-28*: 14; *Lhum-52*: 3; *Lhum-11*: 7; *Lhum_39*: 8; *Lhum_35*: 18; *Lhum-19*: 11; *LihuT1*: 11). To ensure the quality of the markers, the sample size, the number of alleles and effective alleles, and the observed and expected heterozygosity were also calculated in each colony and in each population using GenAlEx (Peakall & Smouse, 2006; Table S4). Hardy-Weinberg disequilibrium were tested in each colony using Genepop 4.7.4 (Rousset, 2008; Table S4). Linkage disequilibrium was tested between all pairs of loci using Genepop 4.7.4 (Rousset, 2008; Table S5). These analyses confirmed the overall good quality of the primers. The genotypes are available on the repository.

The mean relatedness between the workers’ genotypes was calculated using the method of Queller & Goodnight (1989). This allowed us to determine the genotypic similarity of microsatellite markers between pairs of nestmate individuals compared to an expected value between two individuals taken at random from the population. Negative values indicate that the degree of kinship between the two individuals tested was less than that of individuals drawn randomly from the population. Based on these values, the averaged relatedness between nestmate workers have been calculated within each colony. We also calculated the number of private alleles within each colony.

Finally, to get an estimation of the relationship among the studied colonies, we first calculated the *F_st_* between all pairs of colonies as a relative measure of genetic differentiation given the total genetic variation present in the population throughout the sampling range (GenAlEx software; Peakall & Smouse, 2006). We also determined the number of genetically homogeneous groups using the Bayesian clustering algorithm implemented in the software Structure v. 2.3.1 (Pritchard et al., 2010) based on the admixture model with correlated allele frequencies.

We used two models, one without apriori regarding the structure and one with a prior on the colony (Hubisz *et al*., 2009). The number of a priori unknown clusters (K) varied from K = 1 to 12 (*i.e*., the number of colonies), with 10 iteration runs for each K-value. Each run consisted of 500,000 replicates of the Markov Chain Monte Carlo after a burn-in of 500,000 replicates. The 10 independent runs were analysed using CLUMPAK (Kopelman *et al*., 2015) to group sets of highly similar runs into modes and generate a consensus solution for each of them, and we kept the majority mode (*i.e.*, the mode grouping the largest number of similar runs). CLUMPAK was used to define the optimal K-value using the deltaK method of Evanno et al. (2005) and the mean of the logarithm of the data probabilities per K (mean lnP(K)) were calculated using Structure Harvester (Earl and vonHoldt, 2012).

We further combined two approaches: a direct approach using the LnP(K), and a hierarchical approach following Balkenhol et al. (2014). In the direct approach, we simply considered the optimal number of genetic groups to be the number defined by LnP(K). In the hierarchical approach, we assumed that genetic substructure could occur, and we kept the optimal number of clusters suggested by deltaK to create new groups of individuals. Individuals were grouped assuming a membership coefficient of at least 50% to belong to a cluster (Balkenhol *et al*., 2014). The same process was then separately iterated within each cluster to measure sub-structuring within the identified clusters. All parameters remained identical except for the maximum number of clusters tested, which corresponded to the number of colonies involved in the hierarchical level.

## Results

### Sampling

Sampling success was high in Galiza and Catalunya (6 out of 7 locations; 85.71%), finding *L. humile* in three Galizan islands and four mainland locations, *i.e*. in all visited locations except in O Facho, a Galizan mainland area where these ants were not previously detected (Table S1). Despite the higher effort performed (Table S1; Figure S2), sampling success was lower in Greece (44.45%), only finding *L. humile* in two populations of one island (Crete) and in two populations of the mainland (Table 1; see also this table for population codes). As a previously reported study showed two distinct genetic clusters in Galiza (Sanmartín-Villar, *et al*., 2022), we sampled an additional colony (“Refoxos”; REF) to estimate the southern boundary extent of the northern cluster. To simplify the island mainland combination pairs, it was excluded from other analyses.

### Morphometrics

Ants showed different values for all the three parameters (eyespan, face width and face height) measured between locations (island or mainland), region (Galiza or Greece) and the parameter interaction with region or locality (Table 2, Figure 2). Ants from the Cretan island (HRK and HSS) and Catalunya showed the lowest values in comparison with all the other colonies (t> 7.42, *p*< 0.001), and Cretan and Catalunyan ants showed similar size (t< 1.42, *p*> 0.626). No differences were found in any of the three measurements taken between ants from the island and mainland populations of Galiza (t< 0.92, *p*> 0.918). Ants from Galiza showed similar eye span and face width to the mainland populations from Greece (t< 1.90, *p*> 0.231; Figure 2A-B), but they have smaller face height (t> 0.263, *p*< 0.026; Figure 2C).

**Figure 1.**
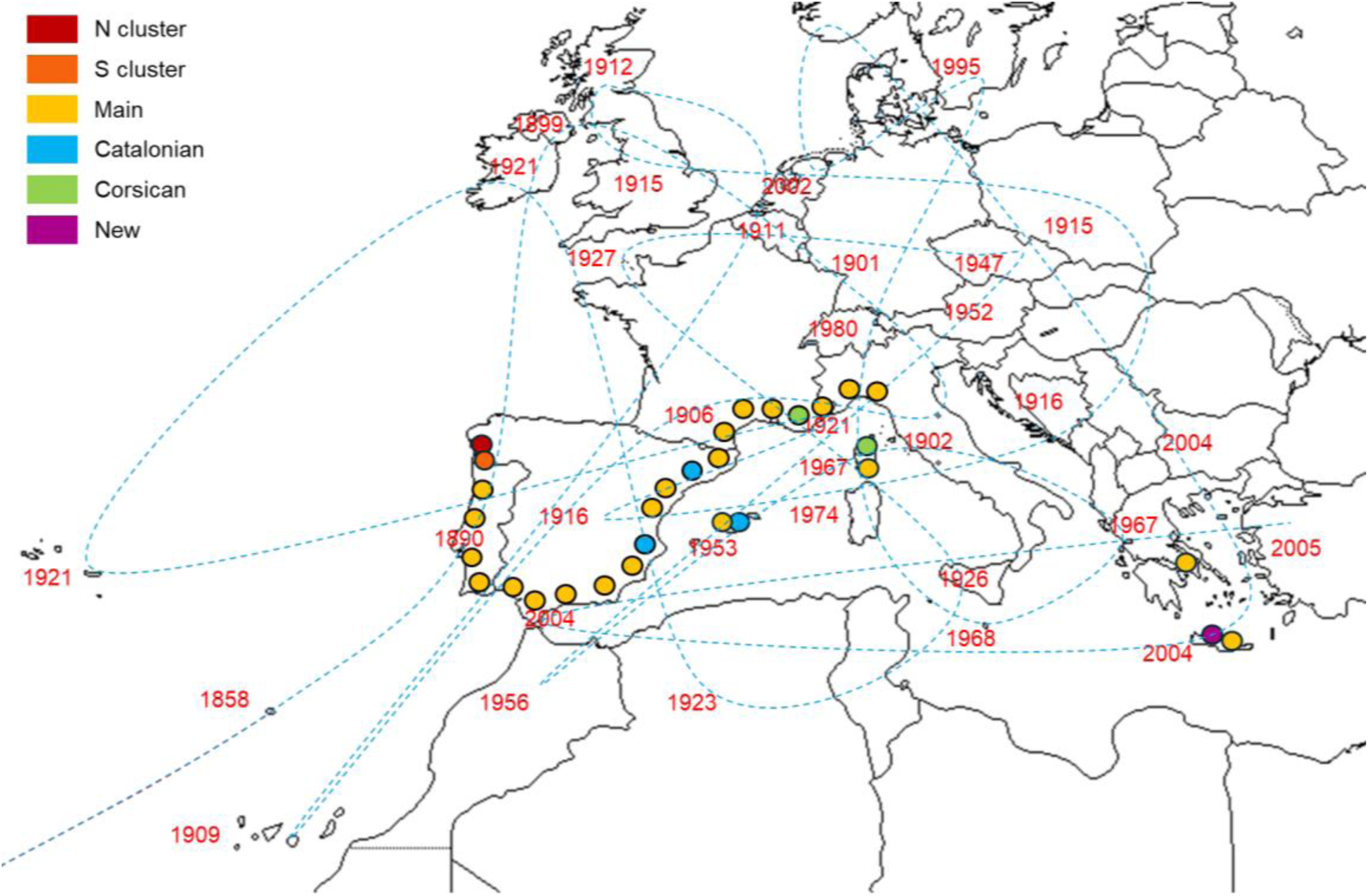
Theoretical colonisation route of the Argentine ant (blue dashed line) in Europe and North Africa, using the introduction in each region as the first detection date (red). When ants were detected in different regions in the same year, the nearest region was taken as the next region. The complexity of the lines stresses the difficulty of understanding the past distribution of the Argentine ant and the need for improvement. Based on Wetterer et al. 2009. Circles summarise the location of the published population diversity. Yellow: main supercolony (Giraud et al. 2002). Blue: Catalunyan supercolony (Giraud et al. 2002, pers. comm. Xavier Espadaler 2021); Green: Corsican supercolony (Blight et al. 2010). Orange and maroon: Galizan clusters (Sanmartín-Villar 2022). Violet: Heraklian supercolony (this work).

**Figure 2.**
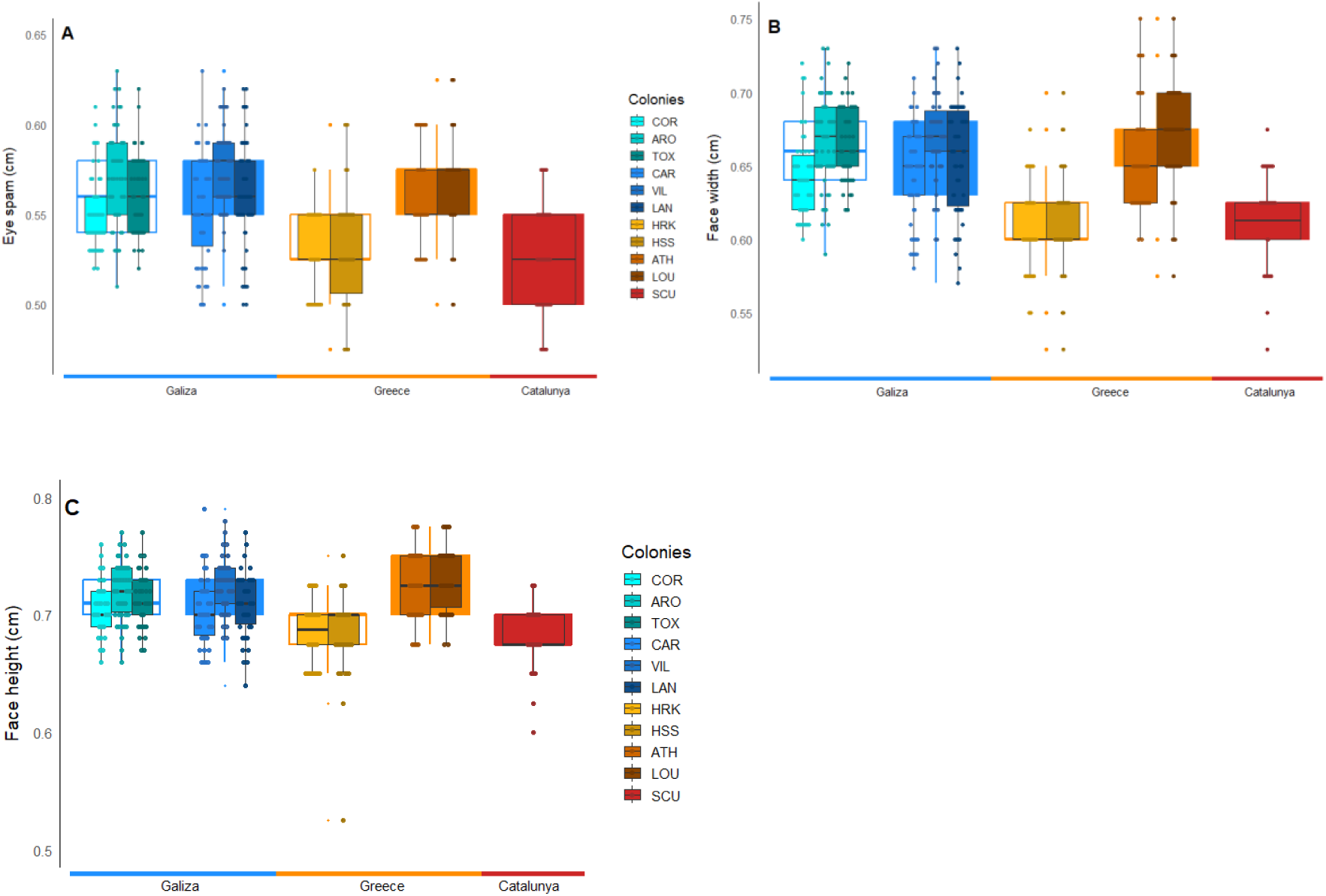
Results of the morphometric analyses showing differences between combination kinds and combinations for maximum eyes-span (**A**), face height (**B**), face width (**C**). Empty boxes: Islands. Filled boxes: Mainland.

**Table 2.** Comparison of morphometric measurements between ants of different populations.

| **Variable** | **Comparison** | **Value** | **DF;F** | ***p*** |
| --- | --- | --- | --- | --- |
| **Eyespan** | Location: island-mainland | 14.03 | 1;545 | <0.001 |
|  | Region: Galiza-Greece | 65.05 | 1;545 | <0.001 |
|  | Location X Region | 40.41 | 1;545 | <0.001 |
| **Face width** | Location: island-mainland | 19.23 | 1;545 | <0.001 |
|  | Region: Galiza-Greece | 72.56 | 1;545 | <0.001 |
|  | Location X Region | 93.81 | 1;545 | <0.001 |
| **Face height** | Location: island-mainland | 19.17 | 1;545 | <0.001 |
|  | Region: Galiza-Greece | 45.58 | 1;545 | <0.001 |
|  | Location X Region | 69.94 | 1;545 | <0.001 |

### Agonism tests

With the exception of a single case (COR-CAR; 1 out of 630), no biting behavior was observed in Galiza. In Greece, interactions involving Heraklion ants showed lethal interactions involving biting in 84 out of 90 cases with 16 cases of mortality, while no biting was recorded in any other Greek interactions (0 out of 210).

The number and the duration of the agonistic interactions differed between the kind of combination (Table 3, Figure 3A-B). The number and duration of the agonistic interactions were higher among Greek populations (1.33±0.08 interactions; 70.56±8.52 s) than among Galizan ones (0.38±0.03 interactions; 4.27±0.72 s; number: t = -10.85, p < 0.001; duration: t = -7.75, p < 0.001).

**Figure 3.**
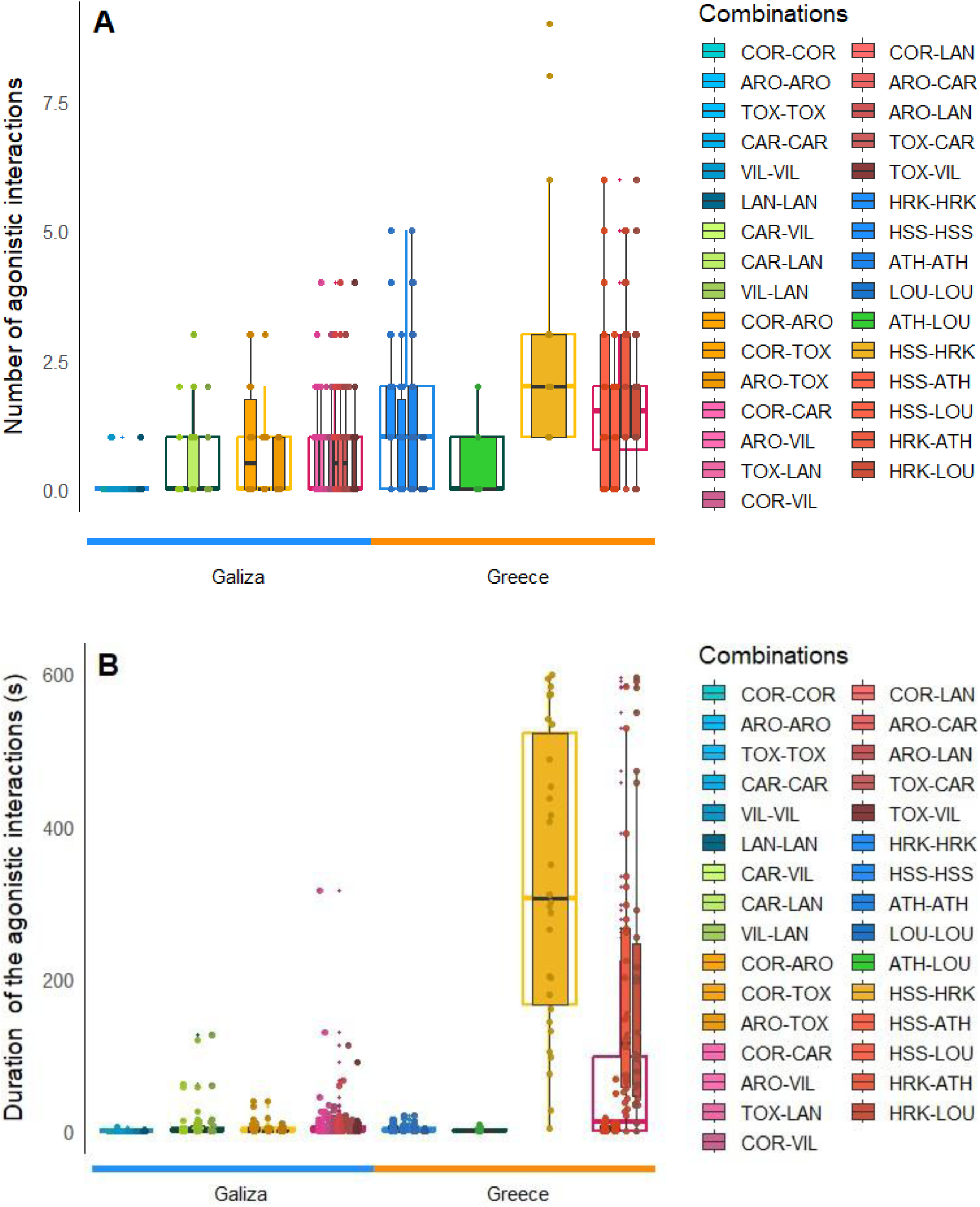
Results of the agonism tests showing the number of agonistic interactions by combination kind and for each combination (**A**); and the duration of the agonistic interactions by combination type and for each combination (**B**).

**Table 3.**
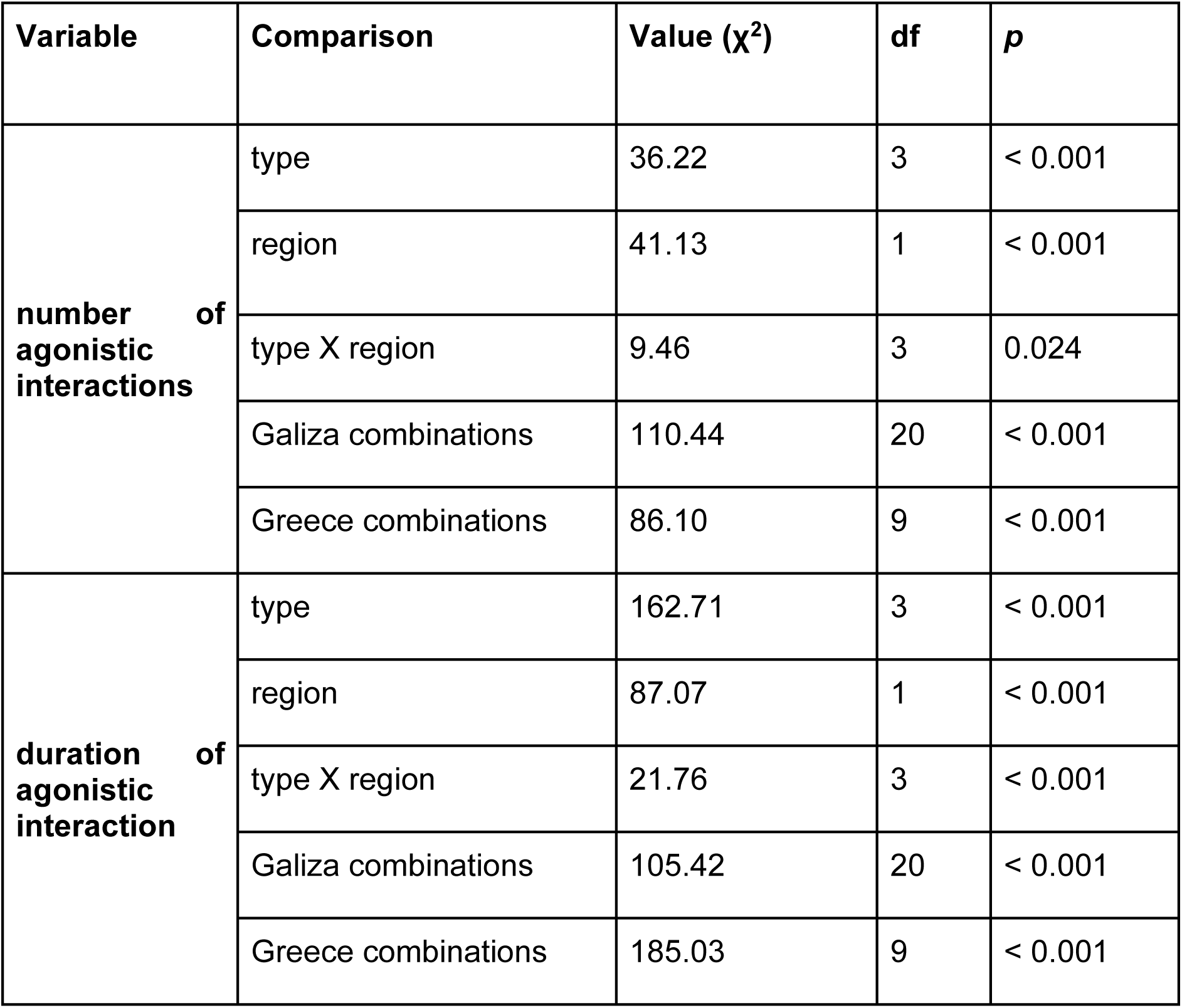
Comparison of agonistic interaction parameters between populations.

| Variable | Comparison | Value ($\chi^2$) | df | *p* |
| --- | --- | --- | --- | --- |
| **number of agonistic interactions** | type | 36.22 | 3 | < 0.001 |
|  | region | 41.13 | 1 | < 0.001 |
|  | type X region | 9.46 | 3 | 0.024 |
|  | Galiza combinations | 110.44 | 20 | < 0.001 |
|  | Greece combinations | 86.10 | 9 | < 0.001 |
| **duration of agonistic interaction** | type | 162.71 | 3 | < 0.001 |
|  | region | 87.07 | 1 | < 0.001 |
|  | type X region | 21.76 | 3 | < 0.001 |
|  | Galiza combinations | 105.42 | 20 | < 0.001 |
|  | Greece combinations | 185.03 | 9 | < 0.001 |

A higher number of agonistic interactions than control combinations (control vs islands: Z= -4.28, *p*< 0.001; control vs island-mainland: Z= -6.33, *p*< 0.001) were found in Galiza, but no differences were found for the rest of the combinations (island-island and mainland-mainland) in this region (Z< 1.87, *p*> 0.246). No differences were found when comparing the number of antagonistic interactions in the combinations performed with Greek colonies (Z< 1.50, *p*> 0.824).

The duration of the agonistic interactions in ants from Galiza was higher only in combinations when comparing to the control groups (control vs mainland-mainland: Z= -5.07, *p*< 0.001; control vs island-island: Z= -6.15, *p*< 0.001; control vs island-mainland: Z= -10.42, *p*< 0.001), while no differences were found in the duration of the agonistic interactions for other combinations (Z< 1.45, *p*> 0.442). The duration of the agonistic interactions differed in most combination kinds in Greece (control and island-island: Z= -5.76, *p*< 0.001; control and island-mainland: Z= -6.65, *p*< 0.001; island-island and mainland-mainland: Z: 4.83, *p*< 0.001; island-mainland and mainland-mainland: Z= 5.05, *p*< 0.001) but not in the comparison between control and mainland-mainland populations and between island-island and island-mainland populations (Z< 1.55, *p*> 0.240).

### Cuticular hydrocarbon analysis

The CHC profiles of *Linepithema humile* populations included a total of 23 distinct peaks, corresponding to 41 different hydrocarbons when co-eluting compounds were considered. These compounds can be grouped into three major categories: straight-chain alkanes, methyl-branched alkanes, and alkenes. Seven straight-chain alkanes were detected (n-C23 to n-C32), together with 17 methyl-branched alkanes (3 mono-, 12 di-, and 11 trimethyl alkanes), which represented the largest class in the profiles. In addition, six monoenes ranging from C18:1 to C28:1 were present. The last three peaks, two trimethyl and one dimethyl alkanes, together accounted for nearly 50% of the total profile. While most hydrocarbons were detected in all populations, a few showed restricted distributions (Table S3).

The NMDS ordination reveals strong geographic structuring of CHC profiles, while also highlighting noteworthy within-area differentiation (Figure 4). This pattern is confirmed by the global PERMANOVA, which detected highly significant differences among populations (R²= 0.917, F = 24.3, *p*< 0.001). Galizan samples form the largest cluster, with most localities overlapping tightly, suggesting high similarity in their CHC profiles. However, CHC samples extracted from ants collected in TOX and COR islands are separated from the main Galizan cluster (R²= 0.684, F= 34.6, *p*= 0.0001). The samples from mainland Greece (ATH, LOU) tend to overlap in the ordination, indicating that their CHC profiles are highly similar. By contrast, Crete does not form a cohesive unit, with CHC samples from HRK occupying the extreme left of the NMDS space, whereas those from HSS plotted more centrally, between Galizan and Greek mainland. The NMDS thus indicates marked differences between the two Cretan localities, consistent with the high proportion of variance explained in the pairwise PERMANOVA (R²= 0.950, F= 76.3, p= 0.1), although the limited sample size likely reduced statistical power.

**Figure 4.**
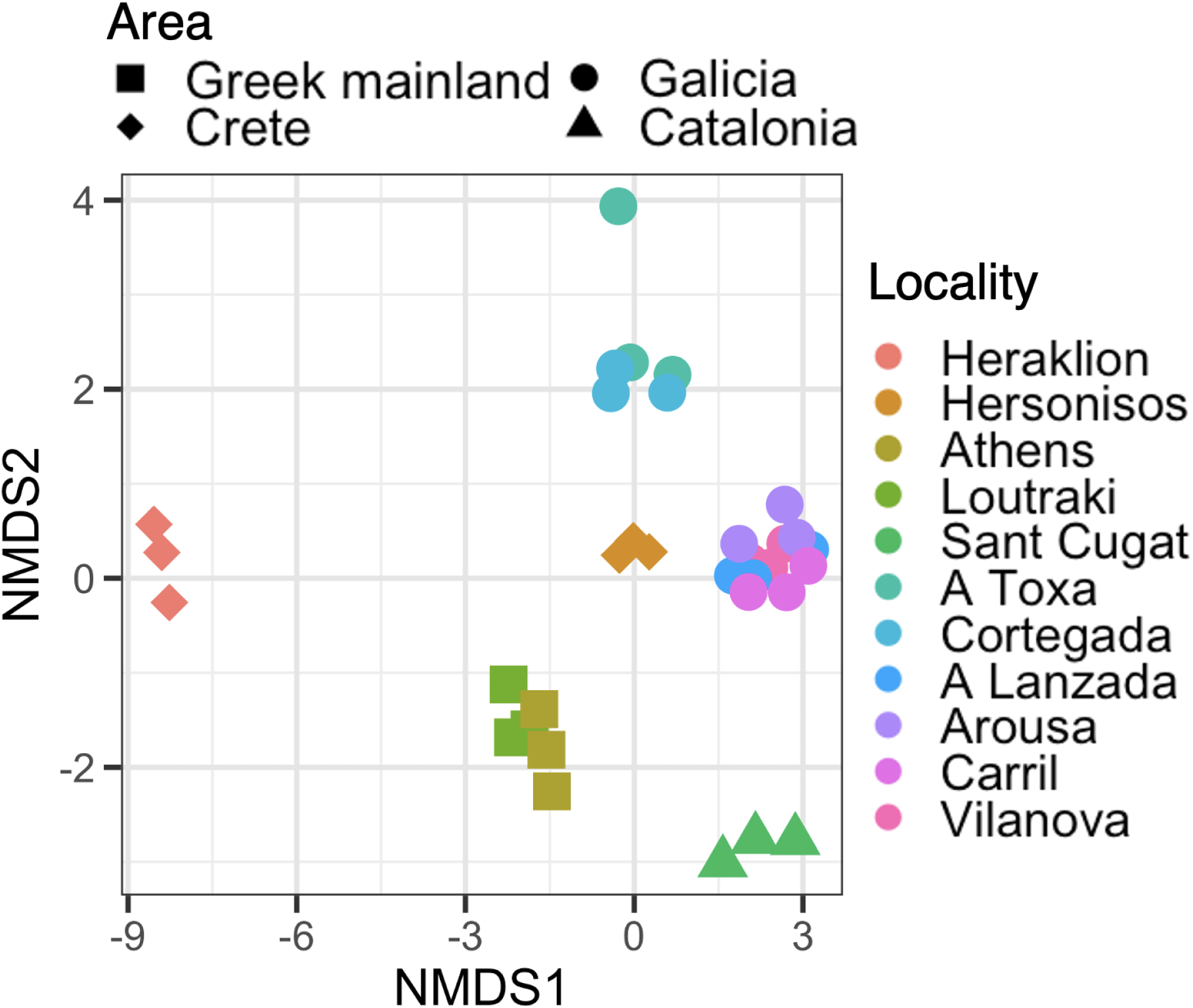
Non-metric multidimensional scaling (NMDS) plot based on Euclidean distances among chemical profiles of *L. humile* workers, calculated from log-transformed absolute abundances of CHC peaks (STRESS=0.07).

Distance-based analyses highlight the unexpected pattern mentioned: despite being geographically part of Crete, the CHC profiles of HSS samples are significantly closer to those of the main Galizan cluster than to either HRK or the Greek mainland (Wilcoxon rank-sum tests: W= 0.0, p< 0.001; W= 68, p< 0.001, respectively).

### Genetic variability

The total number of alleles in each colony ranged between 24 and 32, irrespective to the location or region (number of alleles: ATH (Greece): 24; ARO (Galiza): 25; SCU (Catalunya): 25; LAN (Galiza): 26; LOU (Greece): 27; TOX (Galiza): 28; VIL (Galiza): 29; HSS (Greece): 30; HRK (Greece): 31; COR (Galiza): 31; CAR (Galiza): 31). Three of the four Greek colonies (LOU, HSS, ATH) did not exhibit any private alleles. In Galiza, the number of private alleles per colony ranged from 0 to 2 (ARO: 0, VIL: 0, COR: 1, LAN: 1, CAR: 2, TOX: 2). Only two colonies showed a notably high number of private alleles: the Catalunyan colony (SCU: 11 private alleles) and one Greek colony (HRK: 7 private alleles). These two colonies also ranked among those with the highest nestmate relatedness (Table S7). Additionally, two other colonies (LOU in Greece and ARO in Galiza) showed relatively high within-colony relatedness (>0.4, Table S7). No consistent pattern was found linking these results to whether colonies were located on islands or the mainland.

Pairwise Fst values between colonies suggest that colonies from Greece and Galiza are not genetically isolated from one another (Table 5). At the local scale, mainland colonies are genetically very similar, both in Galiza and Greece (Table 5). Island colonies are not consistently more genetically isolated, either from mainland colonies or from other islands (Table 5). In both geographic regions, however, one island colony appears notably more genetically isolated, regardless of geographic distance from other colonies, with higher genetic differentiation (ARO in Galiza, HRK in Greece; Table 5).

**Table 5.** Pairwise Matrix of *F_st_* Values between all pairs of colonies. *F_st_* values below the grey empty diagonal cells. Very low *F_st_*: 0 - 0.049 (light orange); Low *F_st_*: 0.050 - 0.149 (orange); Medium *F_st_*: 0.150 - 0.250 (red); High *F_st_*: >0.250 (brown). The probability (P(rand >= data)) based on 999 permutations is shown above the grey empty diagonal cells. * : colonies collected in islands. Locations are ordered according to the number of each *F_st_* category explained here with colours.

|  | Galiza<br>CAR | Galiza<br>COR* | Gali<br>za<br>LAN | Greec<br>e LOU | Galiz<br>a<br>TOX<br>* | Greec<br>e<br>HSS* | Greec<br>e ATH | Galiz<br>a VIL | Galiz<br>a<br>ARO<br>* | Greec<br>e<br>HRK* | Catalun<br>ya SCU |
| --- | --- | --- | --- | --- | --- | --- | --- | --- | --- | --- | --- |
| Galiza<br>CAR |  | 0.001 | 0.00<br>1 | 0.001 | 0.00<br>1 | 0.001 | 0.001 | 0.00<br>1 | 0.00<br>1 | 0.001 | 0.001 |
| Galiza<br>COR* | 0.062 |  | 0.00<br>1 | 0.001 | 0.00<br>1 | 0.001 | 0.001 | 0.00<br>1 | 0.00<br>1 | 0.001 | 0.001 |
| Galiza<br>LAN | 0.055 | 0.047 |  | 0.009 | 0.00<br>1 | 0.002 | 0.001 | 0.00<br>1 | 0.00<br>1 | 0.001 | 0.001 |
| Greece<br>LOU | 0.087 | 0.053 | 0.02<br>5 |  | 0.00<br>2 | 0.183 | 0.005 | 0.00<br>1 | 0.00<br>1 | 0.001 | 0.001 |
| Galiza<br>TOX* | 0.074 | 0.061 | 0.02<br>7 | 0.03 |  | 0.001 | 0.001 | 0.00<br>1 | 0.00<br>1 | 0.001 | 0.001 |
| Greece<br>HSS* | 0.098 | 0.056 | 0.03<br>6 | 0.016 | 0.04<br>5 |  | 0.704 | 0.00<br>1 | 0.00<br>1 | 0.001 | 0.001 |
| Greece<br>ATH | 0.107 | 0.068 | 0.04<br>5 | 0.031 | 0.06<br>5 | 0.01 |  | 0.00<br>1 | 0.00<br>1 | 0.001 | 0.001 |
| Galiza<br>VIL | 0.043 | 0.069 | 0.08<br>9 | 0.122 | 0.11<br>7 | 0.133 | 0.151 |  | 0.00<br>1 | 0.001 | 0.001 |
| Galiza<br>ARO* | 0.122 | 0.159 | 0.09<br>5 | 0.149 | 0.09<br>1 | 0.165 | 0.132 | 0.18<br>3 |  | 0.001 | 0.001 |
| Greece<br>HRK* | 0.246 | 0.249 | 0.28<br>6 | 0.316 | 0.27<br>7 | 0.295 | 0.265 | 0.30<br>7 | 0.31<br>4 |  | 0.001 |
| Catalun<br>ya SCU | 0.283 | 0.298 | 0.32<br>8 | 0.366 | 0.34<br>1 | 0.343 | 0.321 | 0.35 | 0.39<br>4 | 0.296 |  |

Both hierarchical and non-hierarchical genetic clustering analyses yielded broadly consistent results (Figure S3), and the models with and without prior information converged. Therefore, we present here the direct models generated with and without prior colony information (see Figure S2 for full results). Six genetically distinct groups were identified (Figure 5), with colonies SCU (Catalunya) and HRK (Greece) standing out as clearly distinct entities. Colony ARO (Galiza) was also well differentiated, confirming the Fst-based findings. A general genetic identity corresponding to Greek or Galizan origin emerged, although colonies LAN and TOX (Galiza) showed intermediate profiles. Colony COR (Galiza) also displayed a distinct profile in the model without prior information, consistent with the hierarchical approach identifying seven clusters (see Figure S2). In conclusion, aside from the reference colony SCU (Catalunya), the three colonies showing particularly distinct genetic profiles are all island colonies (HRK in Greece, ARO and COR in Galiza; Figure 5).

**Figure 5.**
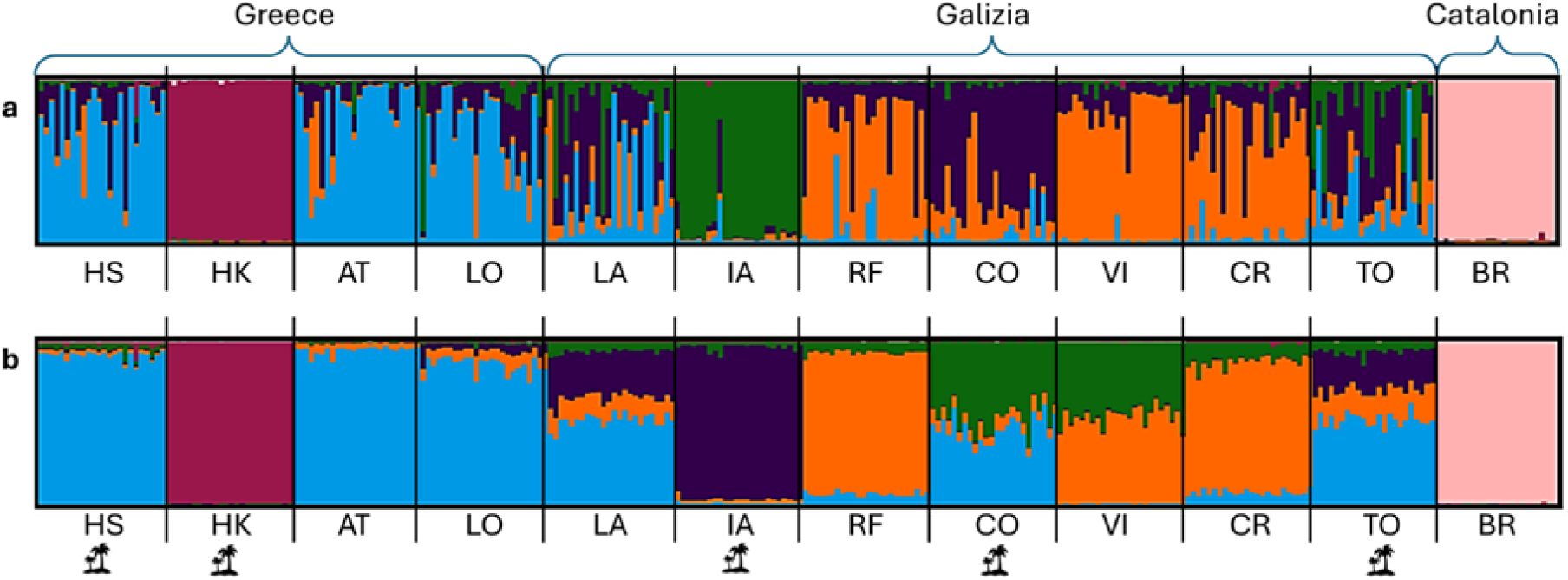
Barplot of Bayesian clustering results without prior information (a) and with prior information based on colony membership (b). In both cases, the estimated number of genetically distinct groups is six (LnP(K) = 6). Sample size: n = 286 individuals.

## Discussion

While *Linepithema humile* was detected across the previously known Galizan locations, its presence in Greece was comparatively scarce, with records restricted to two locations on one of the four sampled islands and two isolated mainland areas. The parameters studied (morphometry, behavior, CHC and genetic profile) varied in different patterns among populations, not showing any unique effect of the assumed allopatry and finding unexpected similarities among workers from Galiza and Greece between islands and mainland populations. Argentine ants from Crete and Catalunya were the smallest, where the ants of the other colonies did not differ in size. Lethal agonism was observed primarily in confrontations involving ants collected in Heraklion, with one exceptional case recorded in Galiza. Argentine ants from the Catalunyan supercolony and those collected in Heraklion showed the most different hydrocarbon composition with the rest, but they also differ between them. Small differences in CHC profile between mainland and islands from Galiza and Greece were also detected. Surprisingly, Hersonissos (Crete) Argentine ants showed a more similar chemical profile with the Argentine ants from Galizan populations than to mainland Greece populations. The microsatellite variability of the ants from the Catalunyan supercolony and those collected in Heraklion did not match with those found in the rest of the colonies, nor did they match with each other. Our results suggest variability within the main supercolony but lower than the expected between supercolonies (Giraud et. al., 2002; Thomas et. al., 2005). Although Cretan ants showed similar size, one population of them (Heraklion), showed clear differences in the three other measured traits; agonistic behaviour, cuticular hydrocarbon profile and genetic differentiation, highlighting the existence of a previously non-identified supercolony in Europe.

Morphology is generally considered a readily identifiable signal for divergence because it is a quantifiable trait of how selection and genetic change alter organismal forms, taking account of ecological and functional differences among populations and species (Zelditch et al., 2012). Studies on other taxonomic clades show an effect of island biogeography on the morphological features (Foster, 1964) in several animal species. We did not see a global trend of gigantism or dwarfism in islands in our study. In fact, few studies have previously reported insular biogeography effects on invertebrates morphology (snails; Cornejo et al., 2022; beetles; Palmer, 2002, de Cerqueira et al., 2023; moths; Yama et al., 2019). The morphological divergence of eusocial insects is shown mainly due to the complex ecology of the region (Gibb et al., 2015; Sosiak & Barden, 2021; Y. Zhao et al., 2020). Morphological measurements of Argentine ants from Galizan islands and mainland did not show a significant difference, but we found some of the trends described before in the Galizan colonies (Fraga-Cimadevila et al., 2025). Samples collected from different islands show substantial morphological differences. Samples from Cortegada and A Toxa islands and the closest mainland samples (Carril and Lanzada) were almost identical in size, whereas the Cretan ants (Heraklion and Hersonissos) were smaller compared to the mainland from Greece. Galiza’s ants were obtained from different island populations geographically close to the mainland, while in Greece the island populations belonged to one island located 300 km apart from mainland. The particular environmental and ecological factors of the Cretan island (temperature, seasonality, productivity, predators and competitors; Clémencet & Doums, 2007; Wills et al., 2014) may have a more pronounced effect on workers’ morphology (Purcell et al., 2016) than the large genetic distance between them and other populations. Evidence from Korea (Jung et al., 2023) and California in North America (Whyte et al., 2023) suggest that the size of the ants are conserved even between different supercolonies albeit not geographically distant. Therefore, morphology seems inaccurate in predicting the divergence of a supercolony, at least when considering a time span such as that elapsed since the introduction of Argentine ants.

The duration and the number of agonistic interactions within Galizan and Greek colonies were higher than control combinations, indicating agonism within the supercolony. However, the intensity of these interactions remained substantially lower than those typically observed between established supercolonies, where biting, carrying, and lethal encounters are common (Giraud et al., 2002; Thomas et al., 2006). The number of agonistic interactions between colonies involving islands were higher than between mainland populations and control. One possible explanation is that ants required repeated inspection of cuticular cues to assess relatedness, leading to increased antennation and low-level agonism (Eyer et al., 2021). These patterns are consistent with local differentiation emerging between island and mainland populations, potentially reflecting early stages of fragmentation within a mosaic supercolony structure.

Our findings support the presence of the main supercolony (Giraud, 2002) in South Eastern Europe, though we found chemical and genetic differentiation that could match with the presence of different clusters (see Sanmartín-Villar, et al., 2022). Galizan colonies showed a chemical clustering that could be a consequence of the colonization process, the proximity of the sampled colonies, and/or similar environmental and dietary constrictions (Menzel, Schmitt and Blaimer, 2017). The similarity between the cuticular profile of Hersonisos (Greece) and Galizan ants suggests a common origin of their introduction or the effect of convergence due to similar local conditions (see for example: Liu et al., 2025; Rodrigues Méndez et al., 2024). Colonies from mainland Greece having different chemical profiles that Hersonissos suggest a different route of invasion, likely from the mainland from other European countries or from a later period. Because CHC composition is responsible for nestmate recognition, variations among populations can influence agonism, connectivity between colonies, and ultimately the spread and resilience of invasive colonies (Buellesbach *et al*., 2018; Holze, Schrader and Buellesbach, 2021; Whyte *et al*., 2023). Genetic distances between colonies suggest that gene flow is generally more frequent on the mainland, although island colonies are not (or at least, not all) genetically isolated from the mainland or from each other. Genetic movement between the islands and corresponding mainland populations might occur actively due to the presence of land bridges or passively through human movements. Our results support that supercolonies are not rigid distinct units, but rather a mosaic of behaviourally defined networks of colonies dependent on genetic and chemical divergences (see Hakala et al., 2020; Seppä et al., 2012).

The genetic and chemical differences found in Cortegada and A Toxa Galizan islands suggest that these populations are exposed to other selection pressures arising from microhabitat shifts (Hais et al., 2024) or lack of individual flow with the mainland colonies. The genetic distance found between Illa de Arousa and Cortegada relative to the mainland colonies showed, respectively, the exact same value as the one found between the Galizan genetic clusters (Fst= 0.13; Sanmartín-Villar, Silva, et al., 2022) and between the main and the Corsican supercolony (Fst= 0.06; Blight et al., 2012). It should also be noted that the Galizan colonies studied by us might be exclusively from the described northern cluster in Galiza (Sanmartín-Villar, Silva, et al., 2022), except the colony from Illa de Arousa, that could belong to the southern cluster or be potentially a separate genetic cluster in the region. Although we found chemical and genetic divergence of these two island colonies from the mainland colonies, the level of agonism remained lower than expected from previous studies with similar genetic distance between them (Blight et al., 2012; Sanmartín-Villar et al., 2022) and from different seasons (see Supplementary Material).

The chemical differentiation between the mainland and the island colonies was higher than the genetic distance, suggesting the faster effect of local ecology (Buczkowski et al., 2005; Liang & Silverman, 2000, but see Mothapo & Wossler, 2016) in comparison with the effect involved in genetic differentiation due to fixation of different alleles under complete isolation (Peña-Carrillo et al., 2021; Sprenger et al., 2021). Variation among allopatric Galizan island and mainland colonies may have emerged primarily in chemical traits, which subsequently influenced nestmate recognition. This divergence likely led to the identification of foreign ants as non-nestmates and triggered agonism. Even when agonism is low, it may prevent colony fusion and ultimately restrict gene flow, promoting genetic isolation. Mapping this variation onto colonization pathways, as in Greek–Galizan comparison, provides a powerful framework to infer past, current and future introductions and spread.

A genetic relatedness level close to that expected under a monogynous-monandrous system has been evidenced in the Catalunyan colony, suggesting either a lower number of queens or a different spatial organization of queens. The location (island-mainland) does not appear to influence genetic relatedness in the colonies studied and thus, structural changes in colony organization seem to not depend on allopatry (see Table S7). However, colonies collected in Galiza tend to consist of less-related workers, suggesting that colony structure might be influenced by broader geographic location, with potentially distinct social systems across the sampled regions. This highlights the social structure plasticity within this species and may reflect ecological differences between regions (Ingram et al. 2002).

The ants collected in Heraklion exhibited (i) high levels of agonism towards individuals from other colonies that ended in lethal outcomes, as has been observed among supercolonies (e.g., Giraud et al., 2002; Thomas et al., 2005); (ii) distinct profiles in the relative abundance of dimethyl- and trimethylalkanes compared to other colonies here studied and to other supercolonies (e.g., Buellesbach et al., 2018; Greene and Gordon, 2007; Whyte et al., 2023); and (iii) genetic differentiation from both the main supercolony (Fst ≈ 0.284) and the Catalunyan supercolony (Fst = 0.296), similar to the previously described between different supercolonies (Giraud et al., 2002). Therefore, our results support the existence of a supercolony hitherto unidentified in Europe, rather than a local differentiation within the main supercolony, which we term the ‘Heraklian supercolony’ in reference to the location of its first discovery. Future studies should focus on unravelling the origin of this potential new supercolony in Europe and its similarity with other supercolonies present in other continents as part of a preventive approach to avoid further introductions into Europe and other continents and explore the mechanisms and consequences of the rapid evolutionary changes in the Argentine ant.

Our data reveals the substantial heterogeneity within the main supercolony in Europe comparable to the previously reported divergence between Corsican and the main supercolony. Heraklian supercolony meets the behavioral, genetic and chemical criteria for a previously unknown supercolony in Europe. The within supercolony variability found restructures the concept of supercoloniality and, with the discovery of a potential new supercolony after more than 150 years of invasions, highlights the lack of knowledge we still have about an invasive species that has a worldwide drastic ecological and economic impact. Due to the diversity of ecological factors driving local population traits, geographic isolation (allopatry) does not reliably predict, at least at this period of time, how populations within a supercolony will diverge. The sample size of colonies found was probably small to predict a definite causative reason for the mosaic-like colony structure previously theorised. Strengthening efforts to study metapopulation dynamics in invasive species integrating CHCs, genetics, behaviour, as well as ecological factors across native and introduced ranges will be crucial to understand how invasion routes, local adaptation and ongoing ecological pressures generate and maintain the diversity observed within globally successful species such as *L. humile* (Blight et al., 2010; Brandt et al., 2009b; Mothapo & Wossler, 2011; Suarez et al., 2001; Torres et al., 2007). Thus a coordinated international research that transcends fieldwork limitations, methodological differences, and the apparent inconsistency of population-level traits is crucial for understanding the evolutionary dynamics of *Linepithema humile*.

## Acknowledgements

Many thanks to Sebastian Salata, Xavier Espadaler, María Dolores Rodríguez Poza, and Stephen Clifford for sharing or posting their observations of *L. humile*. Thanks to Mario Crespo Montalbán, Violette Chiara, Xoan Sanmartín Pazos, and Miguel Barros Nogueira for their assistance during fieldwork performed in Catalunya and Galiza. We are grateful to Adolfo Cordero-Rivera for providing access to the laboratory facilities and equipment used in this study when performing experiments in Galiza. We would also like to thank Gema Trigos-Peral for verifying the species identification of *Linepithema humile* samples from Greece, Daniel Sánchez-García for supporting the CHC extraction methodology and Monika Patrzyk for developing the genetic extraction protocol. This research is part of the Polonez Bis project No. 2021/43/P/NZ8/03306 co-funded by the National Science Centre (Poland) and the European Union Framework Programme for Research and Innovation Horizon 2020 under the Marie Skłodowska-Curie grant agreement no. 945339. E.Cs. was supported by the project No. 2022/45/P/NZ8/04018 cofunded by the National Science Centre and the European Union Framework Programme for Research and Innovation Horizon 2020 under the Marie Skłodowska-Curie grant agreement No. 945339.

