## Supplementary data for "Isolation and introductions disrupt the homogeneity of Argentine ants in Europe"

*co-last authors

**Seasonal effect of agonism**

It has been well established that changes in humidity, temperature and resource availability modulate the behavior of ants according to different seasons (Heller and Gordon, 2006; Gordon, Dektar and Pinter-Wollman, 2013; Gordon *et al.*, 2023; Krapf *et al.*, 2023; Menges *et al.*, 2024). Agonism between the same combination of colonies changes according to the season (Sanmartín-Villar, Cruz Da Silva, *et al.*, 2022; Menges *et al.*, 2024). So to account for the seasonal change in collection between Galiza and Greece, we performed another round of agonism test in September 2025 between the Galizan samples (N=6). The same methodology was followed as before. The number and duration of the agonistic interactions differed among combinations (number: $\chi^2$= 63.73, df= 20, $p$< 0.001; duration: $\chi^2$= 64.50, df= 20, $p$< 0.001). Combinations of ant workers from COR with CAR showed a higher number (z= -3.89, p= 0.0105) and higher duration (z< -3.55, p< 0.0404) of agonistic interactions than control groups. Combinations involving ARO and TOX showed a higher number of interactions than control groups. The duration and total number of interactions were correlated for each combination in both seasons (number of interactions: $\rho(28)$=0.549, p=0.009; duration: $\rho(28)$=0.744, p< 0.001). COR and CAR had similar interactions in both seasons. The agonism interactions correlated across both seasons (Mantel r = 0.445, $p$= 0.044). This result suggests that the seasonal behavioural effect (Heller & Gordon, 2006) may be limited to modifying agonism outcomes only between genetic clusters (Sanmartín-Villar et al., 2022), and that the populations analysed here belong to the North cluster (although see the genetic differentiation detected in ARO). Agonism interactions involving COR and CAR, as well as ARO with VIL (but not TOX or LAN), were interpreted as a *nasty neighbour* effect product of local adaptation and isolation (Fraga-Cimadevila et al., 2025).

Future studies should examine the distribution of genetic clusters in Galiza to better understand within supercolony interactions in this region.

**Table S1.** Sampling effort. *Areas sampled*: Number of areas sampled in the same locality. *Sampled (km)*: kilometres sampled. *Time (h)*: hours spent sampling. The effort performed in Galiza was limited to visiting known populations (no sampled distances or times).

| L. humile | RegionArea | Locality | Acronym | Type | Areas sampled | Sampled (km) |
|---|---|---|---|---|---|---|
| Present | Galiza | Carril | CAR | Mainland | 1 | - |
| Present | Galiza | Cortegada | COR | Island | 1 | - |
| Present | Galiza | Vilanova | VIL | Mainland | 1 | - |
| Present | Galiza | Illa de Arousa | ARO | Island | 1 | - |
| Present | Galiza | Toxa | TOX | Island | 1 | - |
| Absent | Galiza | O Facho | - | Mainland | 1 | - |
| Present | Galiza | Lanzada | LAN | Mainland | 1 | - |
| Present | Galiza | Refoxos | REF | Mainland | 1 | - |
| Present | Catalunya | Sant Cugat | BR | Mainland | 1 | - |
| Present | Greece | Athens | ATH | Mainland | 3 | 6.6 |
| Present | Greece | Loutraki | LOU | Mainland | 2 | 2.7 |
| Absent | Greece | Piraeus | - | Mainland | 1 | 2.2 |
| Absent | Greece | Corinthos | - | Mainland | 1 | 11.2 |
| Absent | Greece | Aegina | - | Island | 12 | 6.4 |
| Absent | Greece | Milos | - | Island | 8 | 14.2 |
| Absent | Greece | Santorini | - | Island | 3 | 20.5 |
| Present | Greece | Heraklion | HRK | Island | 6 | 12.1 |

| | | | | | | |
|---|---|---|---|---|---|---|
| | | (Crete) | | | | |
| Present | Greece | Hersoniso s (Crete) | HSS | Island | 7 | 6.2 |

**Table S2**. *Post hoc* results for agonism tests. The number of agonistic interactions is shown above the grey diagonal cells, and the duration is shown below them. In each cell, the Z value appears on top and the *p*-value on the bottom.

| | COR COR | ARO ARO | TOX TOX | CAR CAR | VIL VIL | LAN LAN | CAR VIL | CAR LAN | VIL LAN | COR ARO | COR TOX | ARO TOX | COR CAR | ARO VIL | TOX LAN | COR VIL | COR LAN | ARO CAR | ARO LAN | TOX CAR | TOX VIL |
|---|---|---|---|---|---|---|---|---|---|---|---|---|---|---|---|---|---|---|---|---|---|
| **CORCOR** | | 1.86 1.000 | 0.00 1.000 | 0.00 1.000 | 0.00 1.000 | -0.80 1.000 | 1.66 1.000 | **3.68 0.021** | -1.66 1.000 | **4.58 <0.001** | -1.68 1.000 | 2.65 0.607 | **3.77 0.015** | **3.89 0.010** | -1.80 1.000 | -2.90 0.306 | **-4.37 0.001** | **5.36 0.000** | **4.30 0.002** | -1.93 1.000 | -2.56 0.762 |
| **AROARO** | 1.64 1.000 | | 1.86 1.000 | 1.86 1.000 | 1.86 1.000 | 1.06 1.000 | 0.20 1.000 | -1.82 1.000 | 0.20 1.000 | -2.73 0.513 | 0.18 1.000 | -0.80 1.000 | -1.91 1.000 | -2.03 1.000 | 0.05 1.000 | -1.05 1.000 | -2.51 0.846 | **-3.51 0.040** | -2.44 1.000 | -0.07 1.000 | -0.70 1.000 |
| **TOXTOX** | 0.00 1.000 | 1.64 1.000 | | 0.00 1.000 | 0.00 0.500 | 0.80 1.000 | 1.66 1.000 | **3.68 0.021** | -1.66 1.000 | **4.58 <0.001** | 1.68 1.000 | 2.66 0.603 | **3.77 0.015** | **3.88 0.010** | 1.80 1.000 | 2.90 0.305 | **4.37 0.001** | **5.36 0.000** | **4.30 0.001** | 1.93 1.000 | -2.56 0.756 |
| **CARCAR** | 0.00 1.000 | 1.63 1.000 | 0.00 1.000 | | 0.00 1.000 | -0.80 1.000 | -1.66 1.000 | **-3.68 0.021** | -1.66 1.000 | **-4.59 <0.001** | -1.68 1.000 | 2.66 0.611 | **-3.77 0.015** | **3.89 0.010** | -1.81 1.000 | -2.91 0.308 | **-4.37 0.001** | **5.37 <0.001** | **4.30 0.002** | -1.93 1.000 | -2.56 1.000 |
| **VIL VIL** | 0.00 1.000 | 1.64 1.000 | 0.00 0.500 | 0.00 1.000 | | 0.80 1.000 | 1.66 1.000 | **3.68 0.021** | 1.66 1.000 | **4.58 <0.001** | 1.68 1.000 | 2.66 0.600 | **3.77 0.015** | **3.89 0.010** | 1.80 1.000 | 2.90 0.303 | **4.37 0.01** | **5.36 0.000** | **4.31 0.002** | 1.93 1.000 | 2.56 0.751 |

| | | | | | | | | | | | | | | | | | | | | | |
|---|---|---|---|---|---|---|---|---|---|---|---|---|---|---|---|---|---|---|---|---|---|
| **LANLAN** | -0.79<br>1.000 | 0.98<br>1.000 | 0.66<br>1.000 | -0.66<br>1.000 | 066<br>1.000 | | 0.87<br>1.000 | 2.89<br>0.317 | -0.87<br>1.000 | **3.79<br>0.014** | 0.02<br>1.000 | 1.86<br>1.000 | 2.97<br>0.250 | 3.09<br>0.172 | -1.01<br>1.000 | 2.11<br>1.000 | **3.58<br>0.031** | **4.57<br><0.001** | **3.51<br>0.040** | -1.13<br>1.000 | -1.76<br>1.000 |
| **CARVIL** | 1.79<br>1.000 | -0.15<br>1.000 | 1.79<br>1.000 | -1.78<br>1.000 | 1.79<br>1.000 | 1.13<br>1.000 | | 2.02<br>1.000 | 0.00<br>1.000 | -2.92<br>0.297 | -0.02<br>1.000 | 0.99<br>1.000 | -2.11<br>1.000 | 2.22<br>1.000 | -0.14<br>1.000 | -1.24<br>1.000 | -2.71<br>0.535 | **3.70<br>0.020** | 2.64<br>0.623 | -0.27<br>1.000 | -0.89<br>1.000 |
| **CARLAN** | **3.92<br>0.008** | -2.28<br>1.000 | **3.92<br>0.008** | **-3.92<br>0.008** | **3.92<br>0.008** | 3.26<br>0.098 | 2.12<br>1.000 | | 2.02<br>1.000 | -0.90<br>1.000 | 2.00<br>1.000 | -1.03<br>1.000 | -0.09<br>1.000 | 0.20<br>1.000 | 1.88<br>1.000 | 0.78<br>1.000 | -0.69<br>1.000 | 1.68<br>1.000 | 0.62<br>1.000 | 1.75<br>1.000 | 1.12<br>1.000 |
| **VILLAN** | -1.66<br>1.000 | -0.24<br>1.000 | -1.87<br>1.000 | -1.87<br>1.000 | 1.87<br>1.000 | -1.21<br>1.000 | -0.08<br>1.000 | 2.04<br>1.000 | | 2.92<br>0.295 | 0.02<br>1.000 | 0.99<br>1.000 | 2.11<br>1.000 | 2.23<br>1.000 | 0.14<br>1.000 | 1.24<br>1.000 | 2.71<br>0.532 | **3.70<br>0.020** | 2.64<br>0.618 | 0.26<br>1.000 | 0.89<br>1.000 |
| **CORARO** | **4.20<br>0.002** | -2.57<br>0.798 | **4.20<br>0.002** | **-4.20<br>0.002** | **4.20<br>0.002** | **3.54<br>0.034** | -2.41<br>1.000 | -0.28<br>1.000 | 2.33<br>1.000 | | 2.90<br>0.30 | -1.93<br>1.000 | 0.81<br>1.000 | -0.70<br>1.000 | 2.78<br>0.437 | 1.68<br>1.000 | 0.21<br>1.000 | 0.78<br>1.000 | -0.28<br>1.000 | 2.65<br>0.598 | 2.03<br>1.000 |
| **CORTOX** | -1.68<br>1.000 | -0.01<br>1.000 | 1.64<br>1.000 | -1.64<br>1.000 | 1.64<br>1.000 | 0.98<br>1.000 | 0.15<br>1.000 | 2.27<br>1.000 | -0.23<br>1.000 | 2.56<br>0.805 | | 0.97<br>1.000 | 2.09<br>1.000 | 2.20<br>1.000 | -0.12<br>1.000 | -1.22<br>1.000 | 2.69<br>0.555 | **3.68<br>0.021** | 2.62<br>0.649 | -0.25<br>1.000 | -0.87<br>1.000 |
| **AR** | 2.53 | - | 2.53 | 2.53 | 2.6 | 1.87 | 0.7 | - | 0.65 | - | 0.88 | | -1.12 | -1.23 | 0.85 | -0.25 | -1.72 | 2.71 | 1.65 | 0.73 | 0.10 |

| | | | | | | | | | | | | | | | | | | | | | |
|---|---|---|---|---|---|---|---|---|---|---|---|---|---|---|---|---|---|---|---|---|---|
| **OTOX** | 0.875 | 0.89 1.000 | 0.869 | 0.881 | 5 0.599 | 1.000 | 3 1.000 | 1.39 1.000 | 1.000 | 1.68 1.000 | 1.000 | | 1.000 | 1.000 | 1.000 | 1.000 | 1.000 | 0.530 | 1.000 | 1.000 | 1.000 |
| **CORCAR** | **3.75 0.016** | -2.12 1.000 | **3.75 0.016** | **-3.75 0.016** | **3.75 0.016** | 3.09 0.163 | -1.96 1.000 | 0.16 1.000 | 1.88 1.000 | 0.45 1.000 | 2.11 1.000 | -1.22 1.000 | | 0.11 1.000 | 1.96 1.000 | 0.86 1.000 | -0.60 1.000 | 1.59 1.000 | 0.53 1.000 | 1.84 1.000 | 1.21 1.000 |
| **AROVIL** | **3.88 0.010** | -2.24 1.000 | **3.88 0.010** | **3.88 0.010** | **3.88 0.010** | 3.22 0.111 | 2.08 1.000 | -0.04 1.000 | 2.00 1.000 | -0.32 1.000 | 2.23 1.000 | -1.35 1.000 | 0.12 1.000 | | 2.08 1.000 | 0.98 1.000 | -0.49 1.000 | 1.48 1.000 | 0.42 1.000 | 1.95 1.000 | 1.33 1.000 |
| **TOXLAN** | -1.80 1.000 | -0.23 1.000 | 1.87 1.000 | -1.86 1.000 | 1.86 1.000 | -1.21 1.000 | -0.07 1.000 | 2.05 1.000 | -0.01 1.000 | 2.34 1.000 | -0.22 1.000 | 0.66 1.000 | 1.88 1.000 | 2.01 1.000 | | 1.10 1.000 | 2.57 0.749 | **3.56 0.032** | 2.50 0.879 | 0.12 1.000 | -0.75 1.000 |
| **CORVIL** | -2.90 0.306 | -1.56 1.000 | 3.19 0.119 | -3.19 0.120 | 3.19 0.118 | 2.53 0.866 | -1.40 1.000 | 0.72 1.000 | 1.32 1.000 | 1.01 1.000 | -1.55 1.000 | -0.66 1.000 | 0.56 1.000 | 0.68 1.000 | 1.32 1.000 | | 1.47 1.000 | 2.46 0.971 | 1.40 1.000 | 0.97 1.000 | 0.34 1.000 |
| **CORLAN** | **-4.37 0.001** | -2.67 0.613 | **4.31 0.002** | **-4.31 0.002** | **4.31 0.002** | **3.65 0.023** | -2.52 0.875 | -0.39 1.000 | 2.43 1.000 | -0.10 1.000 | 2.67 0.619 | -1.78 1.000 | -0.55 1.000 | -0.43 1.000 | 2.44 1.000 | 1.12 1.000 | | 0.99 1.000 | -0.07 1.000 | 2.44 1.000 | 1.81 1.000 |

| | | | | | | | | | | | | | | | | | | | | | |
|---|---|---|---|---|---|---|---|---|---|---|---|---|---|---|---|---|---|---|---|---|---|
| **AROCAR** | **5.05<br>0.000** | -3.41<br>0.056 | **5.05<br>0.000** | **5.05<br>0.000** | **5.36<br>0.000** | **4.39<br>0.001** | 3.26<br>0.098 | 1.13<br>1.000 | 3.18<br>0.124 | 0.84<br>1.000 | 3.41<br>0.057 | 2.52<br>0.867 | 1.29<br>1.000 | 1.17<br>1.000 | 3.18<br>0.122 | 1.85<br>1.000 | 0.74<br>1.000 | | 1.06<br>1.000 | 3.44<br>0.051 | 2.81<br>0.405 |
| **AROLAN** | **4.25<br>0.002** | -2.61<br>0.714 | **4.25<br>0.002** | **4.25<br>0.002** | **4.30<br>0.002** | **3.59<br>0.029** | 2.45<br>1.000 | 0.33<br>1.000 | 2.37<br>1.000 | 0.04<br>1.000 | 2.60<br>0.721 | 1.72<br>1.000 | 0.49<br>1.000 | 0.37<br>1.000 | 2.38<br>1.000 | 1.05<br>1.000 | -0.06<br>1.000 | 0.80<br>1.000 | | 2.50<br>0.879 | 1.75<br>1.000 |
| **TOXCAR** | -1.93<br>1.000 | -0.32<br>1.000 | 1.96<br>1.000 | -1.95<br>1.000 | 1.95<br>1.000 | -1.29<br>1.000 | -0.16<br>1.000 | 1.96<br>1.000 | 0.08<br>1.000 | 2.25<br>1.000 | -0.31<br>1.000 | 0.57<br>1.000 | 1.79<br>1.000 | 1.92<br>1.000 | 0.09<br>1.000 | 1.24<br>1.000 | 2.35<br>1.000 | 3.09<br>0.163 | 2.29<br>1.000 | | -0.63<br>1.000 |
| **TOXVIL** | -2.56<br>0.762 | -0.98<br>1.000 | -2.61<br>0.716 | -2.61<br>0.724 | 2.61<br>0.711 | -1.95<br>1.000 | -0.82<br>1.000 | 1.30<br>1.000 | 0.74<br>1.000 | 1.59<br>1.000 | -0.97<br>1.000 | -0.08<br>1.000 | 1.14<br>1.000 | 1.26<br>1.000 | -0.74<br>1.000 | 0.58<br>1.000 | 1.69<br>1.000 | 2.43<br>1.000 | 1.63<br>1.000 | -0.66<br>1.000 | |

**Table S3**. Relative mean abundances (% ± SE) of cuticular hydrocarbon (CHC) peaks detected in *Linepithema humile* workers across localities. Each row corresponds to a CHC peak (CHC1–CHC23), with compound identity indicated when available. Columns report values for each sampled locality, grouped by area (Galicia, Catalunya, Greece mainland and island). Dashes (---) indicate compounds not detected in that locality.

| | | Galicia | | | | | | | | | Catalonia | Peloponese | | Crete | |
|---|---|---|---|---|---|---|---|---|---|---|---|---|---|---|---|
| n | CHC | A Toxa | Cortegada | A Lanzada | Arousa | Carril | Catoira | Chazo | Refoxos | Vilanova | Sant Cugat | Athens | Loutraki | Heraklion | Hersonisos |
| 1 | C18:1 | 0.79 ±0.08 | 1.34 ±0.49 | 1.29 ±0.38 | 0.87 ±0.21 | 1.15 ±0.22 | 0.7 ±0.06 | 0.76 ±0.07 | 0.9 ±0.06 | 0.84 ±0.16 | -- | 1.29 ±0.1 | 1.87 ±0.17 | 0.94 ±0.1 | 0.5 ±0.02 |
| 2 | diMe-C18 | --- | --- | 1.94 ±0.4 | 0.86 ±0.24 | 0.88 ±0.07 | 1.16 ±0.22 | 1.22 ±0.05 | 1.28 ±0.03 | 2.37 ±0.14 | 1.16 ±0.09 | 0.78 ±0.15 | 1.52 ±0.3 | 2.69 ±0.29 | 1.74 ±0.16 |
| 3 | C20:1 | 2.42 ±0.16 | 2.43 ±0.48 | 2.22 ±0.34 | 1.76 ±0.19 | 1.91 ±0.19 | 1.69 ±0.11 | 1.88 ±0.03 | 1.91 ±0.28 | 1.72 ±0.13 | 1.83 ±0.43 | 2.52 ±0.34 | 2.73 ±0.06 | 3.54 ±0.23 | 2.17 ±0.24 |
| 4 | C22:1 | 2.26 ±0.06 | 2.28 ±0.37 | 1.8 ±0.19 | 1.56 ±0.19 | 1.48 ±0.1 | 1.37 ±0.05 | 1.51 ±0.13 | 1.77 ±0.08 | 1.43 ±0.16 | 1.78 ±0.17 | 2.24 ±0.25 | 2.19 ±0.1 | 3.44 ±0.35 | 2.03 ±0.09 |
| 5 | triMeC22+n-C23 | 7.24 ±0.28 | 4.01 ±0.85 | 4.97 ±0.25 | 4.91 ±0.29 | 5.64 ±0.55 | 5.53 ±0.28 | 5.32 ±0.36 | 3.12 ±0.7 | 4.7 ±0.26 | 4.82 ±0.91 | 3.21 1.11 | 2.29 ±0.07 | 9.04 ±0.52 | 6.17 ±0.43 |
| 6 | C24:1 | 1.61 ±0.06 | 1.58 ±0.26 | 1.23 ±0.23 | 1.01 ±0.15 | 1.31 ±0.23 | 1.18 ±0.13 | 1.69 ±0.2 | 1.26 ±0.07 | 1.13 ±0.05 | 1.43 ±0.64 | 1.57 ±0.11 | 1.86 ±0.16 | 2.46 ±0.14 | 1.26 ±0.12 |
| 7 | triMeC24+n-C25 | 1.33 ±0.08 | 2.11 ±0.42 | 1.21 ±0.07 | 1.08 ±0.07 | 1.75 ±0.32 | 1.85 ±0.04 | 1.65 ±0.07 | 1 ±0.13 | 1.21 ±0.18 | 2.57 ±1.14 | 1.72 ±0.13 | 1.28 ±0.16 | 2.69 ±0.67 | 2.43 ±0.21 |
| 8 | unknown | 5.91 ±0.19 | 4.23 ±0.77 | 3.29 ±0.33 | 2.73 ±0.29 | 4.17 ±0.25 | 3.91 ±0.23 | 3.36 ±0.41 | 2.21 ±0.17 | 2.15 ±0.4 | 4.43 ±0.72 | 4.69 ±0.35 | 3.63 ±0.23 | 3.81 ±0.83 | 3.89 ±0.68 |
| 9 | C26:1 | 0.93 ±0.17 | 0.8 ±0.18 | 0.86 ±0.1 | 1.07 ±0.2 | 1.23 ±0.16 | 0.86 ±0.05 | 1.28 ±0.11 | 1.13 ±0.08 | 0.89 ±0.05 | 1.59 ±0.57 | 0.94 ±0.05 | 0.87 ±0.23 | 1.44 ±0.06 | 1.12 ±0.11 |
| 10 | triMe-C25+n-C26 | 0.88 ±0.45 | 1.78 ±0.33 | 1.28 ±0.23 | 1.23 ±0.2 | 1.49 ±0.13 | 1.43 ±0.11 | 1.92 ±0.1 | 2.08 ±0.07 | 1.91 ±0.15 | 1.68 ±0.3 | 1.25 ±0.07 | 1.22 ±0.03 | --- | 1.99 ±0.22 |
| 11 | unknown | 4.68 ±0.38 | 3.34 ±0.21 | 4.65 ±0.26 | 4.33 ±0.54 | 3.9 ±0.27 | 4.52 ±0.17 | 5.28 ±0.31 | 6.54 ±0.56 | 4.82 ±0.44 | 3.65 ±0.48 | 4.07 ±0.23 | 6.66 ±0.43 | 4.49 ±0.82 | 4.8 ±0.35 |
| 12 | n-C27 | 5.98 ±0.72 | 7.18 ±0.4 | 5.98 ±0.35 | 5.04 ±0.33 | 4.02 ±0.32 | 5.75 ±0.42 | 5.18 ±0.24 | 4.28 ±0.3 | 7.73 ±0.3 | 7.25 ±3.06 | 6.24 ±0.05 | 6.91 ±0.56 | 3.26 ±0.39 | 18.6 ±0.81 |
| 13 | 3-MeC27 | 3.19 ±0.89 | 1.44 ±0.37 | 1.59 ±0.29 | 1.78 ±0.21 | 1.06 ±0.08 | 1.15 ±0.14 | 1.4 ±0.19 | 2.6 ±0.16 | 2.37 ±0.34 | -- | -- | -- | -- | 0.88 ±0.09 |
| 14 | n-C28+C28:1 | 3.33 ±0.53 | 2.78 ±0.15 | 3.08 ±0.2 | 2.99 ±0.17 | 2.93 ±0.2 | 2.72 ±0.17 | 3.3 ±0.13 | 3.9 ±0.29 | 3.21 ±0.04 | 3.76 ±0.8 | 2.56 ±0.07 | 3.26 ±0.09 | 3.95 ±0.26 | 3.97 ±0.1 |
| 15 | n-C29 | 2.32 ±0.3 | 3.27 ±0.35 | 2.45 ±0.26 | 3.99 ±0.37 | 1.57 ±0.1 | 1.76 ±0.2 | 2.06 ±0.12 | 2.76 ±0.11 | 3.16 ±0.16 | 1.21 ±0.06 | 2.98 ±0.34 | 3 ±0.46 | 5.02 ±0.43 | 7.48 ±0.31 |
| 16 | 3-MeC29 | 1.61 ±0.3 | 2.1 ±0.38 | 1.54 ±0.03 | 2.9 ±0.32 | 1.5 ±0.17 | 1.73 ±0.07 | 1.76 ±0.11 | 2.08 ±0.13 | 2.18 ±0.06 | 1.5 ±0.07 | -- | -- | -- | 1.17 ±0.1 |
| 17 | triMeC29+n-C30 | --- | --- | 1.51 ±0.17 | 1.37 ±0.17 | 1.59 ±0.19 | 1.38 ±0.15 | 1.79 ±0.26 | 1.68 ±0.33 | 1.23 ±0.09 | 1.67 ±0.66 | -- | -- | -- | -- |
| 18 | 3-MeC31+5,9-+5,13-+5,15+5,17-diMeC31 | 1.97 ±0.1 | 2.03 ±0.37 | 1.6 ±0.05 | 2.96 ±0.08 | 1.6 ±0.03 | 1.54 ±0.08 | 1.36 ±0.17 | 1.69 ±0.23 | 1.73 ±0.08 | 1.19 ±0.29 | 1.6 ±0.17 | 1.36 ±0.13 | 2.12 ±0.12 | 1.45 ±0.04 |
| 19 | 5,13,15-+5,13,17-triMeC31+n-C32 | 1.54 ±0.15 | 3.08 ±0.31 | 2.44 ±0.36 | 1.63 ±0.24 | 3.4 ±0.25 | 3.56 ±0.15 | 3.75 ±0.14 | 3.55 ±0.38 | 4.1 ±0.24 | 4.14 ±0.64 | 1.73 ±0.18 | 1.49 ±0.59 | 3.72 ±0.44 | 1.48 ±0.11 |
| 20 | 5,13-+5,15-+5,17-diMeC33 | 1.63 ±0.26 | 2.83 ±0.17 | 2.61 ±0.37 | 3.74 ±0.42 | 2.97 ±0.08 | 2.7 ±0.23 | 3.02 ±0.15 | 3.35 ±0.2 | 3.19 ±0.15 | 4.11 ±0.65 | 5.1 ±0.35 | 2.01 ±0.4 | -- | 1.54 ±0.05 |
| 21 | 5,13,15-+5,13,17-+5,13,19-triMeC33 | 31.8 ±1.35 | 27.5 ±1.84 | 30.9 ±1.1 | 27.2 ±1.05 | 26.3 ±1.06 | 27.6 ±1.08 | 29.5 ±0.8 | 31.1 ±0.59 | 25.9 ±1.38 | 18.4 ±5.73 | 26.8 ±0.82 | 32.7 ±1.21 | 32.6 ±1.23 | 20.9 ±1.46 |
| 22 | 5,13,15-+5,13,17-triMeC35 | 10.1 ±0.55 | 11.3 ±0.62 | 12 ±0.96 | 12.4 ±0.44 | 11.8 ±0.39 | 11.9 ±0.55 | 9.57 ±0.58 | 9.9 ±0.49 | 10.9 ±0.59 | 12.6 ±3.4 | 14 ±0.59 | 13.7 ±0.74 | 14.8 ±0.8 | 10.8 ±0.63 |
| 23 | 14,16-+14,18-+14,20-+14,22-diMeC36 | 8.42 ±1.81 | 12.7 ±1.27 | 9.58 ±1.72 | 12.7 ±0.54 | 16.3 ±1.28 | 14 ±0.3 | 11.4 ±0.76 | 9.98 ±0.94 | 11.2 ±0.82 | 19.3 ±1.43 | 14.7 ±0.41 | 9.44 ±0.75 | -- | 3.68 ±0.74 |

**Table S4**. General description of the eight microsatellite markers used in the study, in each geographic area (**A**) and in each colony (**B**). *N*: Sample Size; *Na*: Number of alleles; *Ne*: Number of effective alleles; *Ho*: Observed Heterozygosity; *He*: Expected heterozygosity; *F*: Fixation index.

**A**

| Geographic area | Locus | N | Na | Ne | Ho | He | F |
|---|---|---|---|---|---|---|---|
| Greece | Lhum-13 | 96 | 6.000 | 4.149 | 0.583 | 0.759 | 0.231 |
| | Lhum-28 | 94 | 8.000 | 2.863 | 0.383 | 0.651 | 0.411 |
| | Lhum-52 | 95 | 3.000 | 2.543 | 0.453 | 0.607 | 0.254 |
| | Lhum-11 | 95 | 6.000 | 2.603 | 0.305 | 0.616 | 0.504 |
| | Lhum-_39 | 89 | 7.000 | 1.781 | 0.202 | 0.439 | 0.539 |
| | Lhum-_35 | 90 | 10.000 | 2.648 | 0.422 | 0.622 | 0.322 |
| | Lhum-19 | 94 | 8.000 | 3.267 | 0.564 | 0.694 | 0.187 |
| | LihuT1 | 94 | 6.000 | 2.191 | 0.160 | 0.544 | 0.706 |
| Galiza | Lhum-13 | 161 | 8.000 | 3.412 | 0.466 | 0.707 | 0.341 |
| | Lhum-28 | 164 | 6.000 | 2.264 | 0.579 | 0.558 | -0.037 |
| | Lhum-52 | 164 | 2.000 | 1.811 | 0.360 | 0.448 | 0.197 |
| | Lhum-11 | 162 | 6.000 | 3.134 | 0.525 | 0.681 | 0.229 |
| | Lhum-39 | 155 | 2.000 | 1.355 | 0.129 | 0.262 | 0.507 |
| | Lhum-35 | 153 | 13.000 | 2.365 | 0.562 | 0.577 | 0.026 |
| | Lhum-19 | 159 | 7.000 | 3.319 | 0.711 | 0.699 | -0.017 |
| | LihuT1 | 159 | 8.000 | 1.746 | 0.453 | 0.427 | -0.060 |
| Catalunya | Lhum-13 | 23 | 1.000 | 1.000 | 0.000 | 0.000 | #N/A |

| | | | | | | | |
|---|---|---|---|---|---|---|---|
| | Lhum-28 | 23 | 5.000 | 3.527 | 0.870 | 0.716 | -0.214 |
| | Lhum-52 | 23 | 2.000 | 1.996 | 0.609 | 0.499 | -0.220 |
| | Lhum-11 | 23 | 3.000 | 1.301 | 0.174 | 0.232 | 0.249 |
| | Lhum-_39 | 19 | 4.000 | 2.756 | 0.789 | 0.637 | -0.239 |
| | Lhum_35 | 19 | 4.000 | 2.569 | 0.842 | 0.611 | -0.379 |
| | Lhum-19 | 22 | 4.000 | 2.021 | 0.591 | 0.505 | -0.170 |
| | LihuT1 | 22 | 2.000 | 1.198 | 0.182 | 0.165 | -0.100 |

**B**

| Colony | | Lhum-13 | Lhum-28 | Lhum-52 | Lhum-11 | Lhum_39 | Lhum_35 | Lhum-19 | LihuT1 |
|---|---|---|---|---|---|---|---|---|---|
| Greece HSS | N | 24 | 24 | 24 | 24 | 21 | 21 | 24 | 24 |
| | Na | 4 | 3 | 2 | 4 | 5 | 6 | 3 | 3 |
| | Ne | 2.776 | 1.707 | 1.800 | 1.587 | 1.282 | 2.901 | 2.133 | 1.682 |
| | Ho | 0.625 | 0.333 | 0.417 | 0.208 | 0.143 | 0.286 | 0.542 | 0.125 |
| | He | 0.640 | 0.414 | 0.444 | 0.370 | 0.220 | 0.655 | 0.531 | 0.405 |
| | F | 0.023 | 0.195 | 0.062 | 0.437 | 0.351 | 0.564 | -0.020 | 0.692 |
| Greece HRK | N | 24 | 22 | 23 | 23 | 20 | 21 | 23 | 23 |
| | Na | 4 | 5 | 2 | 3 | 6 | 5 | 3 | 3 |
| | Ne | 2.021 | 3.195 | 1.910 | 1.679 | 2.857 | 2.617 | 2.351 | 1.426 |
| | Ho | 0.417 | 0.364 | 0.522 | 0.217 | 0.650 | 0.381 | 0.565 | 0.348 |
| | He | 0.505 | 0.687 | 0.476 | 0.405 | 0.650 | 0.618 | 0.575 | 0.299 |

| | | | | | | | | | |
|---|---|---|---|---|---|---|---|---|---|
| | F | 0.175 | 0.471 | -0.095 | 0.463 | 0.000 | 0.383 | 0.016 | -0.165 |
| Greece ATH | N | 24 | 24 | 24 | 24 | 24 | 24 | 24 | 24 |
| | Na | 4 | 2 | 2 | 2 | 2 | 4 | 5 | 3 |
| | Ne | 2.789 | 1.946 | 1.843 | 1.600 | 1.087 | 2.436 | 1.953 | 1.792 |
| | Ho | 0.542 | 0.583 | 0.292 | 0.333 | 0.083 | 0.458 | 0.417 | 0.042 |
| | He | 0.641 | 0.486 | 0.457 | 0.375 | 0.080 | 0.589 | 0.488 | 0.442 |
| | F | 0.156 | -0.200 | 0.362 | 0.111 | -0.043 | 0.222 | 0.146 | 0.906 |
| Greece LOU | N | 24 | 24 | 24 | 24 | 24 | 24 | 23 | 23 |
| | Na | 4 | 3 | 2 | 3 | 1 | 4 | 5 | 5 |
| | Ne | 2.866 | 1.524 | 1.882 | 1.826 | 1.000 | 1.732 | 2.612 | 1.374 |
| | Ho | 0.750 | 0.250 | 0.583 | 0.458 | 0.000 | 0.542 | 0.739 | 0.130 |
| | He | 0.651 | 0.344 | 0.469 | 0.452 | 0.000 | 0.423 | 0.617 | 0.272 |
| | F | -0.152 | 0.273 | -0.244 | -0.013 | N/A | -0.281 | -0.198 | 0.521 |
| Galiza LAN | N | 24 | 23 | 23 | 22 | 22 | 22 | 22 | 22 |
| | Na | 4 | 4 | 2 | 4 | 2 | 5 | 3 | 2 |
| | Ne | 3.191 | 2.455 | 1.830 | 1.840 | 1.046 | 2.114 | 2.711 | 1.365 |
| | Ho | 0.708 | 0.739 | 0.348 | 0.318 | 0.045 | 0.636 | 0.727 | 0.227 |
| | He | 0.687 | 0.593 | 0.454 | 0.457 | 0.044 | 0.527 | 0.631 | 0.268 |
| | F | -0.032 | -0.247 | 0.233 | 0.303 | -0.023 | -0.208 | -0.152 | 0.151 |

| | | | | | | | | | |
|---|---|---|---|---|---|---|---|---|---|
| Galiza ARO | N | 24 | 24 | 24 | 24 | 22 | 19 | 22 | 23 |
| | Na | 2 | 4 | 2 | 5 | 2 | 4 | 4 | 2 |
| | Ne | 1.043 | 2.436 | 1.280 | 2.462 | 1.095 | 1.856 | 2.890 | 1.139 |
| | Ho | 0.042 | 0.792 | 0.250 | 0.667 | 0.091 | 0.579 | 0.682 | 0.130 |
| | He | 0.041 | 0.589 | 0.219 | 0.594 | 0.087 | 0.461 | 0.654 | 0.122 |
| | F | -0.021 | -0.343 | -0.143 | -0.123 | -0.048 | -0.255 | -0.043 | -0.070 |
| Galiza COR | N | 24 | 24 | 24 | 24 | 23 | 24 | 24 | 24 |
| | Na | 5 | 3 | 2 | 4 | 2 | 7 | 5 | 3 |
| | Ne | 2.736 | 1.917 | 2.000 | 1.983 | 1.941 | 2.194 | 3.480 | 2.554 |
| | Ho | 0.625 | 0.583 | 0.417 | 0.458 | 0.217 | 0.542 | 0.833 | 0.750 |
| | He | 0.635 | 0.478 | 0.500 | 0.496 | 0.485 | 0.544 | 0.713 | 0.609 |
| | F | 0.015 | -0.220 | 0.167 | 0.075 | 0.552 | 0.005 | -0.169 | -0.233 |
| Galiza VIL | N | 23 | 23 | 23 | 23 | 23 | 22 | 24 | 24 |
| | Na | 4 | 3 | 2 | 4 | 2 | 6 | 5 | 3 |
| | Ne | 2.108 | 1.715 | 1.189 | 3.235 | 1.910 | 2.588 | 2.600 | 1.510 |
| | Ho | 0.609 | 0.348 | 0.174 | 0.435 | 0.261 | 0.591 | 0.708 | 0.417 |
| | He | 0.526 | 0.417 | 0.159 | 0.691 | 0.476 | 0.614 | 0.615 | 0.338 |
| | F | -0.158 | 0.166 | -0.095 | 0.371 | 0.452 | 0.037 | -0.151 | -0.234 |
| Galiza CAR | N | 22 | 24 | 24 | 24 | 21 | 22 | 24 | 24 |

| | | | | | | | | | |
|---|---|---|---|---|---|---|---|---|---|
| | Na | 4 | 4 | 2 | 4 | 2 | 7 | 5 | 3 |
| | Ne | 2.170 | 2.554 | 1.753 | 2.723 | 1.208 | 2.790 | 2.537 | 1.792 |
| | Ho | 0.364 | 0.667 | 0.458 | 0.708 | 0.190 | 0.455 | 0.667 | 0.583 |
| | He | 0.539 | 0.609 | 0.430 | 0.633 | 0.172 | 0.642 | 0.606 | 0.442 |
| | F | 0.326 | -0.096 | -0.067 | -0.119 | -0.105 | 0.291 | -0.100 | -0.320 |
| Galiza TOX | N | 20 | 22 | 22 | 22 | 20 | 20 | 20 | 20 |
| | Na | 5 | 4 | 2 | 3 | 2 | 2 | 4 | 6 |
| | Ne | 2.151 | 1.921 | 1.963 | 2.344 | 1.161 | 1.406 | 3.089 | 1.717 |
| | Ho | 0.500 | 0.364 | 0.409 | 0.545 | 0.050 | 0.350 | 0.700 | 0.300 |
| | He | 0.535 | 0.479 | 0.491 | 0.573 | 0.139 | 0.289 | 0.676 | 0.418 |
| | F | 0.065 | 0.241 | 0.166 | 0.049 | 0.640 | -0.212 | -0.035 | 0.281 |
| Galizia RF | N | 24 | 24 | 24 | 23 | 24 | 24 | 23 | 22 |
| | Na | 4 | 3 | 2 | 3 | 2 | 7 | 6 | 5 |
| | Ne | 2.157 | 2.039 | 1.969 | 2.315 | 1.043 | 3.008 | 3.255 | 2.205 |
| | Ho | 0.417 | 0.542 | 0.458 | 0.522 | 0.042 | 0.750 | 0.652 | 0.727 |
| | He | 0.536 | 0.510 | 0.492 | 0.568 | 0.041 | 0.668 | 0.693 | 0.546 |
| | F | 0.223 | -0.063 | 0.069 | 0.082 | -0.021 | -0.124 | 0.059 | -0.331 |
| | F | #N/A | -0.214 | -0.220 | 0.249 | -0.239 | -0.379 | -0.170 | -0.100 |

**Table S5.** Hardy-Weinberg equilibrium tests. Estimation of exact P-Values by the Markov chain method, global estimate of FIS over alleles according to Weir and Cockerham (W&C) and Robertson and Hill (R&H).

| Population | Locus | P-value | SE | W&C | R&H |
|---|---|---|---|---|---|
| Greece HSS | Lhum-13 | 0.7194 | 0.0058 | 0.0443 | -0.0053 |
| | Lhum-28 | 0.2176 | 0.0055 | 0.2154 | 0.1053 |
| | Lhum-52 | 1 | 0 | 0.0837 | 0.0856 |
| | Lhum-11 | 0.0441 | 0.0043 | 0.4537 | 0.2028 |
| | Lhum-39 | 0.098 | 0.0089 | 0.3717 | 0.2605 |
| | Lhum-35 | 0.0002 | 0.0002 | 0.5804 | 0.5912 |
| | Lhum-19 | 0.5878 | 0.0039 | 0.0017 | 0.0728 |
| | LihuT | 0.0006 | 0.0003 | 0.7026 | 0.4037 |
| Greece HRK | Lhum-13 | 0.515 | 0.0093 | 0.1958 | 0.0801 |
| | Lhum-28 | 0.0005 | 0.0004 | 0.4886 | 0.5659 |
| | Lhum-52 | 1 | 0 | -0.0732 | -0.0747 |
| | Lhum-11 | 0.0271 | 0.0018 | 0.4799 | 0.2694 |
| | Lhum-39 | 0.8634 | 0.0091 | 0.0256 | 0.0294 |
| | Lhum-35 | 0.0003 | 0.0001 | 0.4041 | 0.3197 |
| | Lhum-19 | 0.7909 | 0.0031 | 0.0387 | -0.0005 |

| | | | | | |
|---|---|---|---|---|---|
| | LihuT | 1 | 0 | -0.1429 | -0.0849 |
| Greece ATH | Lhum-13 | 0.6701 | 0.0068 | 0.1763 | 0.1158 |
| | Lhum-28 | 0.4316 | 0.0027 | -0.1795 | -0.1826 |
| | Lhum-52 | 0.0825 | 0.0018 | 0.3808 | 0.392 |
| | Lhum-11 | 0.6 | 0.0018 | 0.1321 | 0.1353 |
| | Lhum-39 | 1 | 0 | -0.0222 | -0.0227 |
| | Lhum-35 | 0.0147 | 0.0022 | 0.2425 | 0.229 |
| | Lhum-19 | 0.0965 | 0.008 | 0.1667 | 0.0314 |
| | LihuT | 0 | 0 | 0.9094 | 0.5171 |
| Greece LOU | Lhum-13 | 0.4236 | 0.0083 | -0.1311 | -0.0639 |
| | Lhum-28 | 0.1291 | 0.0036 | 0.2923 | 0.1965 |
| | Lhum-52 | 0.3916 | 0.0023 | -0.2243 | -0.228 |
| | Lhum-11 | 1 | 0 | 0.0078 | 0.0086 |
| | Lhum-39 | No information | | | |
| | Lhum-35 | 0.5833 | 0.009 | -0.2616 | -0.1048 |
| | Lhum-19 | 0.9024 | 0.0055 | -0.1761 | -0.0899 |
| | LihuT | 0.0012 | 0.0006 | 0.5368 | 0.2587 |
| Galiza LAN | Lhum-13 | 0.9864 | 0.0008 | -0.0103 | -0.0380 |
| | Lhum-28 | 0.2739 | 0.0063 | -0.2262 | -0.1608 |

| | | | | | |
|---|---|---|---|---|---|
| | Lhum-52 | 0.3537 | 0.0025 | 0.2542 | 0.2614 |
| | Lhum-11 | 0.0680 | 0.0053 | 0.3241 | 0.1204 |
| | Lhum-39 | No information | | | |
| | Lhum-35 | 0.1345 | 0.0096 | -0.1855 | -0.0859 |
| | Lhum-19 | 0.6421 | 0.0038 | -0.1294 | -0.1521 |
| | LihuT | 0.4298 | 0.0020 | 0.1732 | 0.1780 |
| Galiza ARO | Lhum-13 | No information | | | |
| | Lhum-28 | 0.0145 | 0.0020 | -0.3242 | -0.1663 |
| | Lhum-52 | 1.0000 | 0.0000 | -0.1220 | -0.1242 |
| | Lhum-11 | 0.1115 | 0.0088 | -0.1018 | -0.0435 |
| | Lhum-39 | 1.0000 | 0.0000 | -0.0244 | -0.0249 |
| | Lhum-35 | 0.2447 | 0.0068 | -0.2298 | -0.1040 |
| | Lhum-19 | 0.5620 | 0.0058 | -0.0194 | -0.0261 |
| | LihuT | 1.0000 | 0.0000 | -0.0476 | -0.0486 |
| Galiza COR | Lhum-13 | 0.6116 | 0.0100 | 0.0363 | 0.0279 |
| | Lhum-28 | 0.5609 | 0.0047 | -0.1993 | -0.1232 |
| | Lhum-52 | 0.4359 | 0.0025 | 0.1873 | 0.1920 |
| | Lhum-11 | 0.3758 | 0.0094 | 0.0964 | -0.0015 |

| | | | | | |
|---|---|---|---|---|---|
| | Lhum-39 | 0.0099 | 0.0006 | 0.5669 | 0.5869 |
| | Lhum-35 | 0.2612 | 0.0199 | 0.0261 | 0.0181 |
| | Lhum-19 | 0.4042 | 0.0086 | -0.1486 | -0.1190 |
| | LihuT | 0.0008 | 0.0002 | -0.2123 | -0.2226 |
| Galiza VIL | Lhum-13 | 0.8958 | 0.0039 | -0.1365 | -0.1136 |
| | Lhum-28 | 0.1661 | 0.0038 | 0.1871 | 0.0847 |
| | Lhum-52 | 1.0000 | 0.0000 | -0.0732 | -0.0747 |
| | Lhum-11 | 0.0532 | 0.0027 | 0.3897 | 0.2795 |
| | Lhum-39 | 0.0354 | 0.0012 | 0.4699 | 0.4854 |
| | Lhum-35 | 0.3720 | 0.0139 | 0.0602 | 0.0993 |
| | Lhum-19 | 0.7631 | 0.0080 | -0.1301 | -0.0584 |
| | LihuT | 0.6312 | 0.0052 | -0.2137 | -0.1127 |
| Galiza CAR | Lhum-13 | 0.0112 | 0.0019 | 0.3463 | 0.1355 |
| | Lhum-28 | 0.8709 | 0.0040 | -0.0745 | -0.0481 |
| | Lhum-52 | 1.0000 | 0.0000 | -0.0455 | -0.0464 |
| | Lhum-11 | 0.7560 | 0.0055 | -0.0983 | -0.0564 |
| | Lhum-39 | 1.0000 | 0.0000 | -0.0811 | -0.0829 |
| | Lhum-35 | 0.0360 | 0.0066 | 0.3126 | 0.2563 |

| | | | | | |
|---|---|---|---|---|---|
| | Lhum-19 | 0.3432 | 0.0105 | -0.0792 | -0.0420 |
| | LihuT | 0.0477 | 0.0024 | -0.3010 | -0.1795 |
| Galiza TOX | Lhum-13 | 0.2065 | 0.0100 | 0.0909 | 0.1184 |
| | Lhum-28 | 0.0477 | 0.0037 | 0.2632 | 0.4106 |
| | Lhum-52 | 0.4190 | 0.0028 | 0.1888 | 0.1941 |
| | Lhum-11 | 0.8923 | 0.0021 | 0.0718 | 0.0153 |
| | Lhum-39 | 0.0762 | 0.0012 | 0.6545 | 0.6828 |
| | Lhum-35 | 1.0000 | 0.0000 | -0.1875 | -0.1914 |
| | Lhum-19 | 0.8727 | 0.0034 | -0.0095 | -0.0421 |
| | LihuT | 0.1414 | 0.0118 | 0.3049 | 0.2726 |
| Galiza RF | Lhum-13 | 0.2262 | 0.0096 | 0.2434 | 0.0937 |
| | Lhum-28 | 0.8762 | 0.0024 | -0.0418 | -0.0744 |
| | Lhum-52 | 0.6993 | 0.0020 | 0.0899 | 0.0920 |
| | Lhum-11 | 0.7395 | 0.0033 | 0.1036 | 0.1282 |
| | Lhum_39 | No information. | | | |
| | Lhum_35 | 0.9830 | 0.0027 | -0.1025 | -0.0441 |
| | Lhum-19 | 0.1546 | 0.0093 | 0.0808 | 0.0169 |
| | LihuT | 0.1005 | 0.0070 | -0.3099 | -0.1140 |
| | LihuT | 1.0000 | 0.0000 | -0.0769 | -0.0786 |

**Table S6**. Genepop Linkage disequilibrium for each locus pair across all populations (P-value obtained using Fisher's method).

| Locus pair | $Chi^2$ | df | P-Value |
|---|---|---|---|
| Lhum-13 & Lhum-28 | 15.7373 | 22 | 0.8287 |
| Lhum-13 & Lhum-52 | 19.6927 | 22 | 0.6022 |
| Lhum-28 & Lhum-52 | 17.2816 | 24 | 0.8364 |
| Lhum-13 & Lhum-11 | 12.6561 | 22 | 0.9423 |
| Lhum-28 & Lhum-11 | 22.5367 | 24 | 0.5473 |
| Lhum-52 & Lhum-11 | 21.9215 | 24 | 0.5840 |
| Lhum-13 & Lhum_39 | 15.0229 | 20 | 0.7751 |
| Lhum-28 & Lhum_39 | 16.1048 | 22 | 0.8107 |
| Lhum-52 & Lhum_39 | 17.6950 | 22 | 0.7239 |
| Lhum-11 & Lhum_39 | 18.2998 | 22 | 0.6881 |
| Lhum-13 & Lhum_35 | 18.3180 | 22 | 0.6870 |
| Lhum-28 & Lhum_35 | 18.1552 | 24 | 0.7954 |
| Lhum-52 & Lhum_35 | 28.4440 | 24 | 0.2418 |
| Lhum-11 & Lhum_35 | 27.2845 | 24 | 0.2914 |
| Lhum_39 & Lhum_35 | 26.1406 | 22 | 0.2457 |
| Lhum-13 & Lhum-19 | 17.6012 | 22 | 0.7294 |
| Lhum-28 & Lhum-19 | 31.8422 | 24 | 0.1310 |
| Lhum-52 & Lhum-19 | 21.2978 | 24 | 0.6211 |
| Lhum-11 & Lhum-19 | 28.6178 | 24 | 0.2348 |
| Lhum_39 & Lhum-19 | 30.1224 | 22 | 0.1155 |
| Lhum_35 & Lhum-19 | 29.3492 | 24 | 0.2073 |

| | | | |
|---|---|---|---|
| Lhum-13 & LihuT | 18.1510 | 22 | 0.6970 |
| Lhum-28 & LihuT | 25.6148 | 24 | 0.3730 |
| Lhum-52 & LihuT | 38.0460 | 24 | 0.0343 |
| Lhum-11 & LihuT | 19.7414 | 24 | 0.7114 |
| Lhum_39 & LihuT | 6.0320 | 22 | 0.9997 |
| Lhum_35 & LihuT | 22.1465 | 24 | 0.5705 |
| Lhum-19 & LihuT | 30.5284 | 24 | 0.1678 |

**Table S7**. Queller & Goodnight relatedness values within each colony. Colonies are ranked from highest to lowest nestmate relatedness. Colors represent geographic origin (red: Catalunya; green: Galiza; blue: Greece). *: colonies collected in islands.

| | Mean | SE |
|---|---|---|
| Catalunya_SCU | 0.7200 | 0.1211 |
| Galiza_ARO* | 0.6250 | 0.1670 |
| Greece_HRK* | 0.4978 | 0.1773 |
| Greece_LOU | 0.4385 | 0.2262 |
| Greece_ATH | 0.3283 | 0.2545 |
| Galiza_TOX* | 0.3216 | 0.2933 |
| Galiza_VIL | 0.3160 | 0.2431 |
| Galiza_LAN | 0.3158 | 0.2819 |

| | | |
|---|---|---|
| Greece_HSS* | 0.2949 | 0.2559 |
| Galiza_REF | 0.2672 | 0.2509 |
| Galiza_CAR | 0.2526 | 0.2406 |
| Galiza_COR* | 0.1805 | 0.2496 |

**Figure S1.** Results of the agonism tests performed with ants collected in Autumn in Galiza showing the number of agonistic interactions by combination kind and for each combination (**A**); and the duration of the agonistic interactions by combination type and for each combination (**B**).

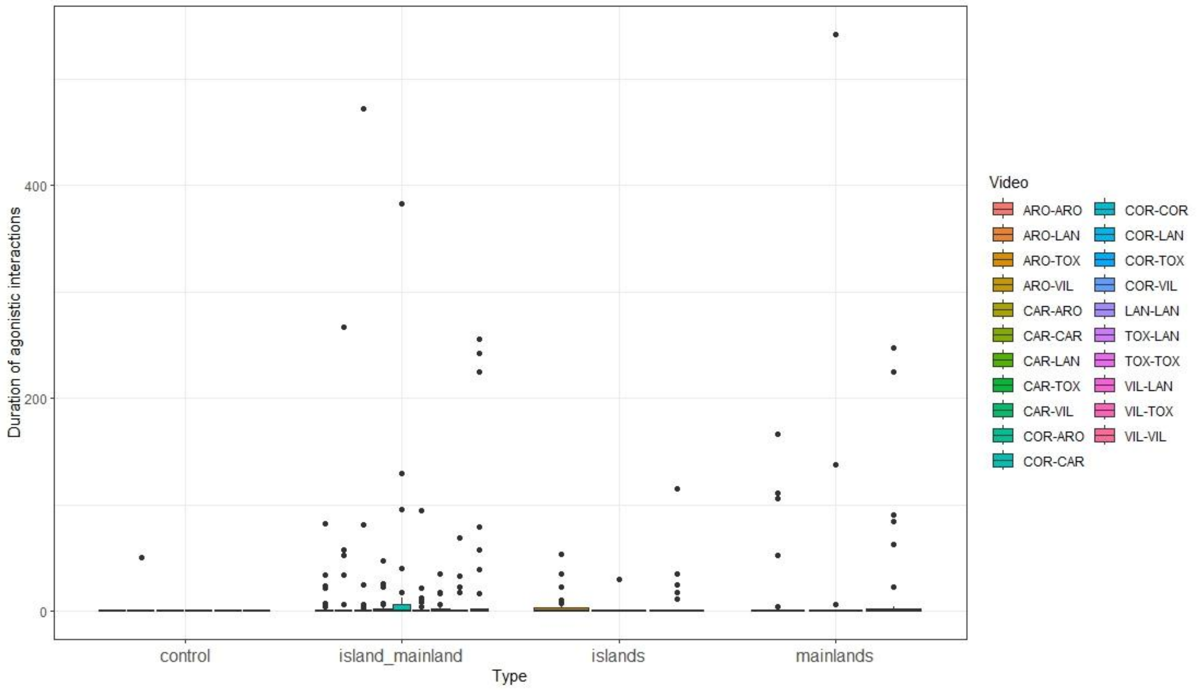

Duration of agonistic interactions
0
200
400
control
island_mainland
islands
mainlands
Type
Video
ARO-ARO
ARO-LAN
ARO-TOX
ARO-VIL
CAR-ARO
CAR-CAR
CAR-LAN
CAR-TOX
CAR-VIL
COR-ARO
COR-CAR
COR-COR
COR-LAN
COR-TOX
COR-VIL
LAN-LAN
TOX-LAN
TOX-TOX
VIL-LAN
VIL-TOX
VIL-VIL


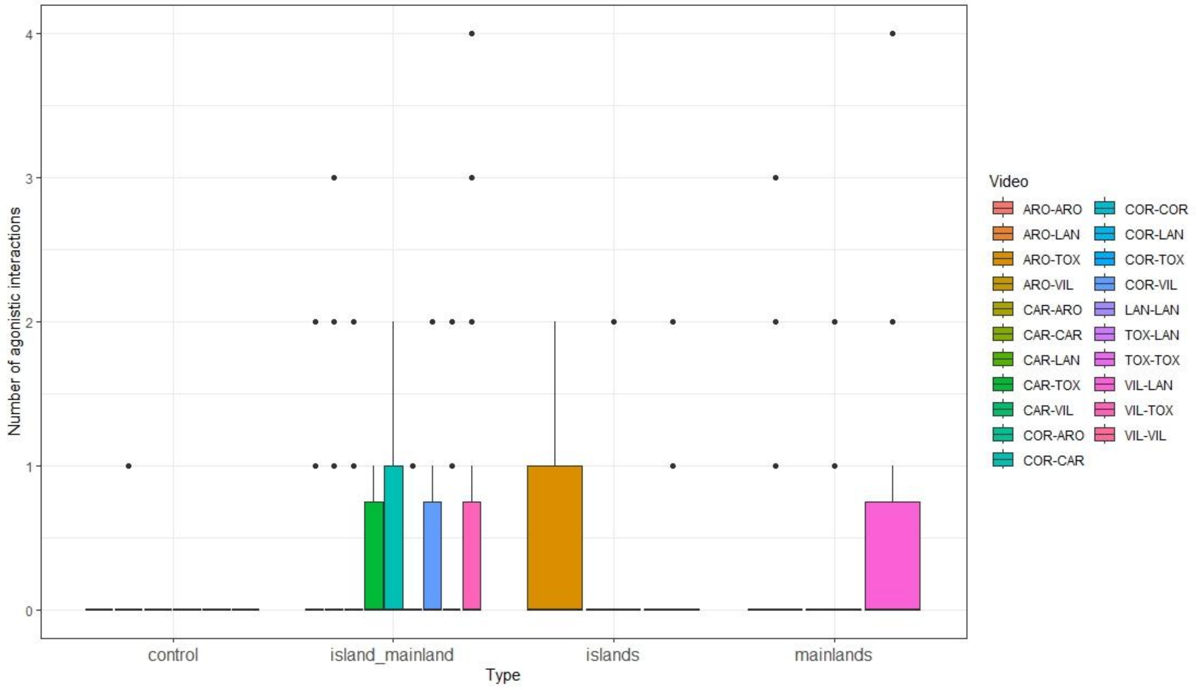

Number of agonistic interactions
0
1
2
3
4
control
island_mainland
islands
mainlands
Type
Video
ARO-ARO
ARO-LAN
ARO-TOX
ARO-VIL
CAR-ARO
CAR-CAR
CAR-LAN
CAR-TOX
CAR-VIL
COR-ARO
COR-CAR
COR-COR
COR-LAN
COR-TOX
COR-VIL
LAN-LAN
TOX-LAN
TOX-TOX
VIL-LAN
VIL-TOX
VIL-VIL

**Figure S2.** Sampled tracks. **A**: Sampled locations in Europe. **B**: Greece. **C**: Galiza. **D**: Athens. **E**: Corinthos and Loutraki. **F**: Aegina. **G**: Milos. **H**: Santorini. **I**: Heraklion. **J**: Hersonissos. Yellow: tracked transects recorded by Google Maps (lines) and locations punctually checked (dots). Red: Locations where the experimental ants were extracted. Black numbers: beeline (black lines) distance (km) within populations. All images are oriented northward. Sant Cugat population (Catalunyan supercolony) omitted.

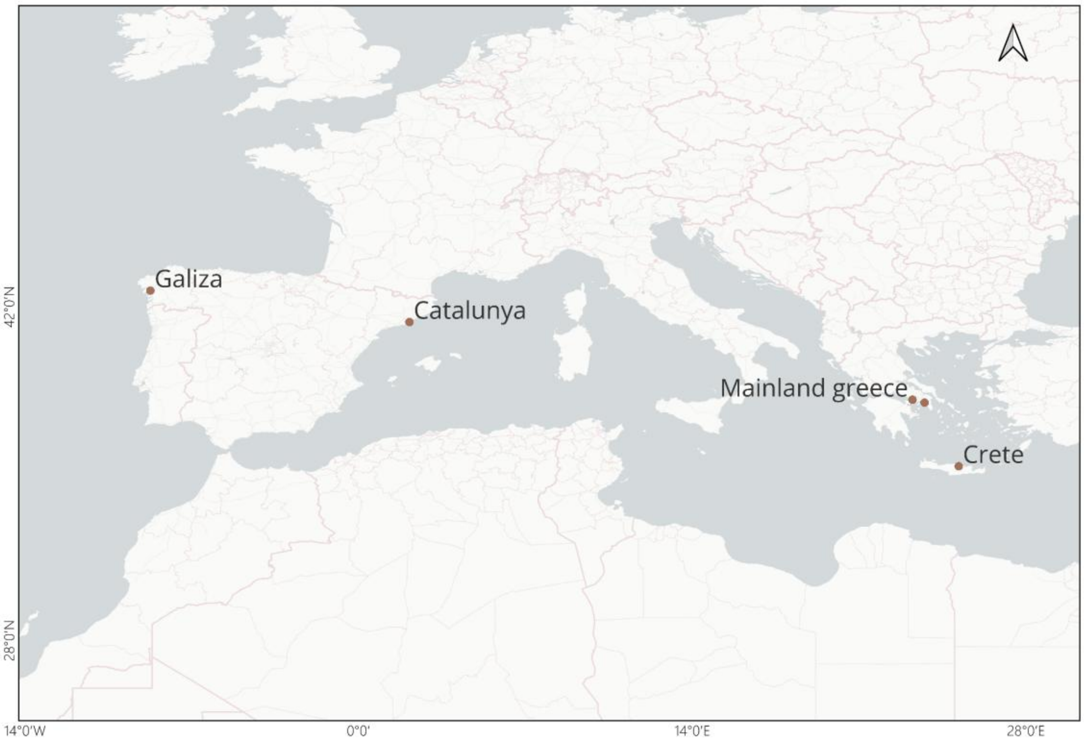
Galiza
Catalunya
Mainland greece
Crete
42°0'N
28°0'N
14°0'W
0°0'
14°0'E
28°0'E

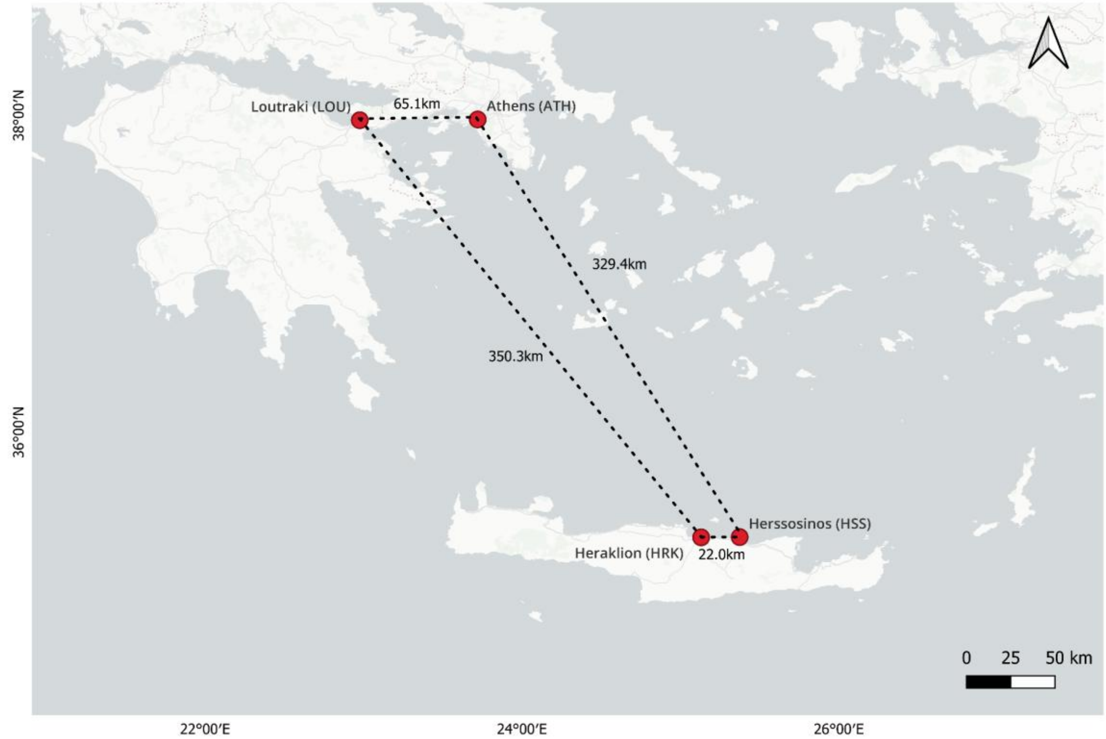
Loutraki (LOU)
65.1km
Athens (ATH)
329.4km
350.3km
Herssosinos (HSS)
Heraklion (HRK)
22.0km
0 25 50 km
38°00'N
36°00'N
22°00'E
24°00'E
26°00'E

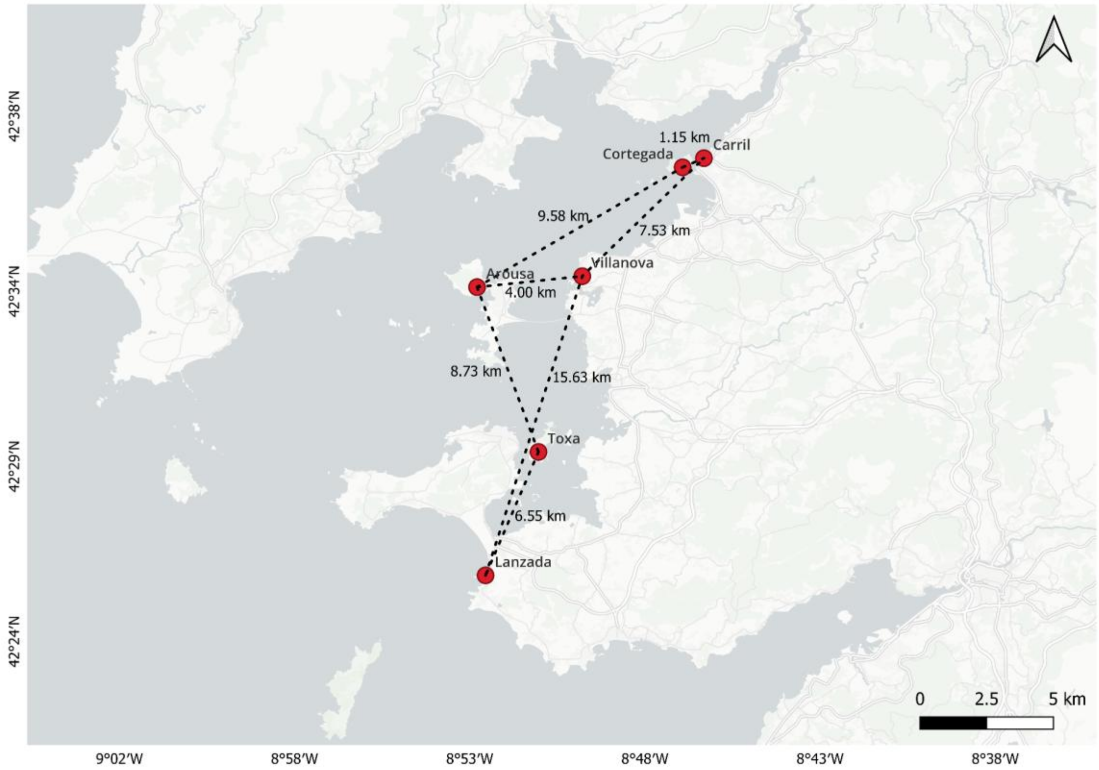
Carril
1.15 km
Cortegada
9.58 km
7.53 km
Arousa
4.00 km
Villanova
8.73 km
15.63 km
Toxa
6.55 km
Lanzada
0
2.5
5 km
42°38′N
42°34′N
42°29′N
42°24′N
9°02′W
8°58′W
8°53′W
8°48′W
8°43′W
8°38′W

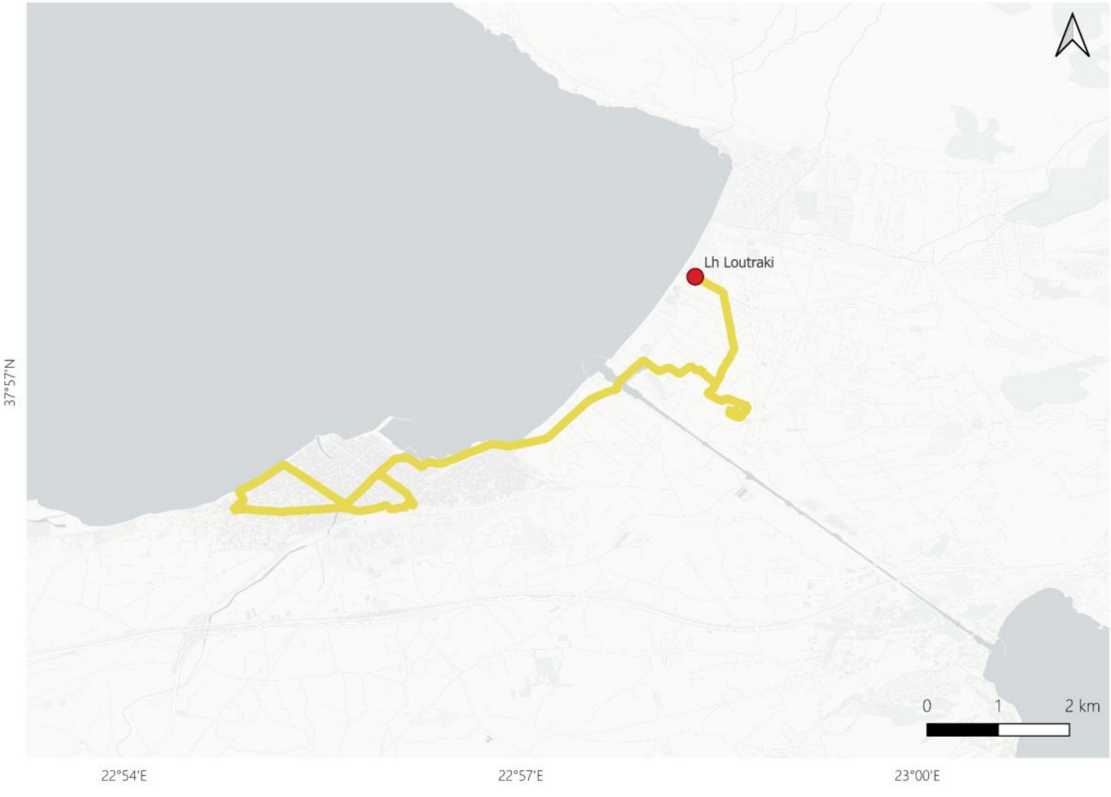
Lh Loutraki
0
1
2 km
37°57′N
22°54′E
22°57′E
23°00′E

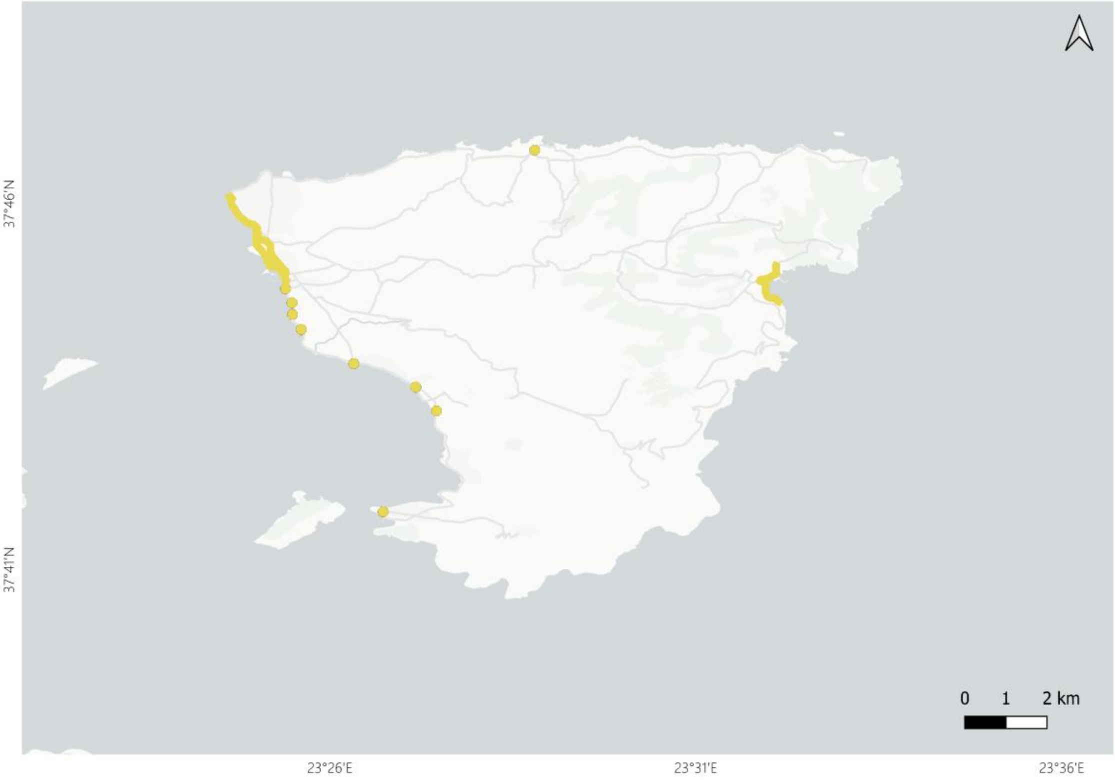
37°46'N
37°41'N
0 1 2 km
23°26'E
23°31'E
23°36'E

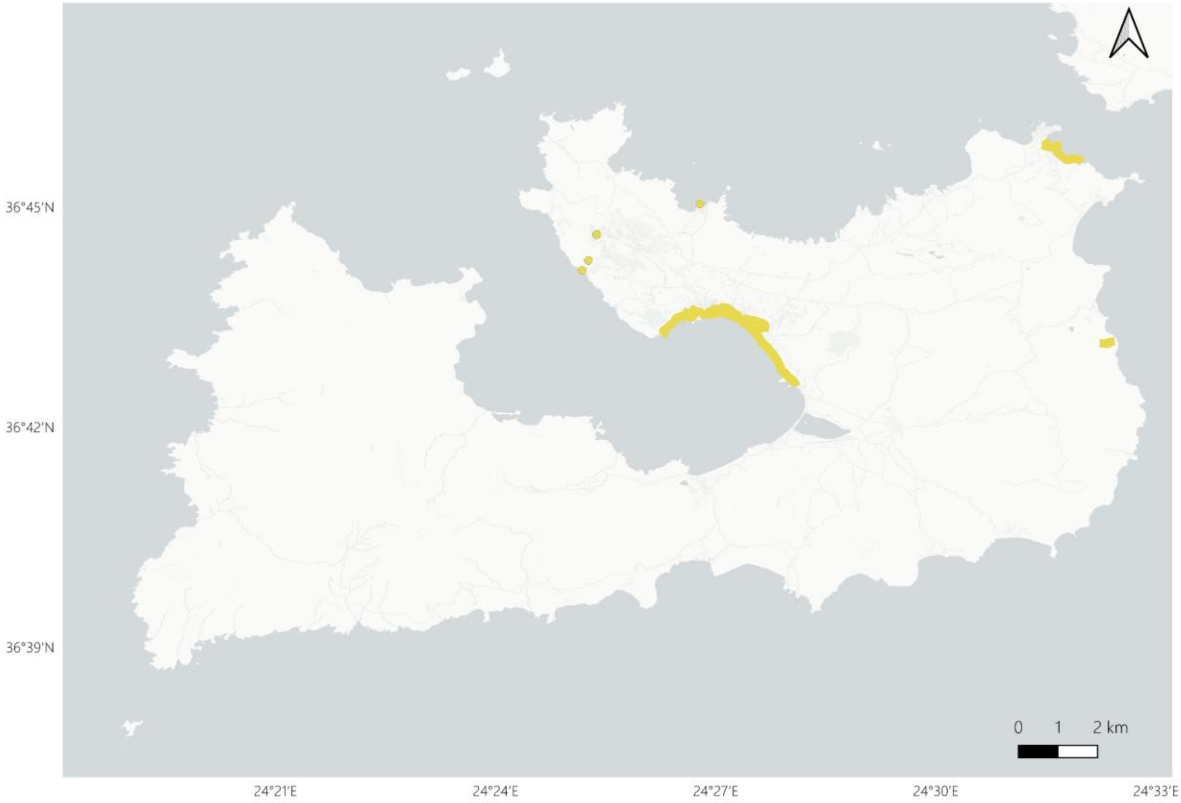
36°45'N
36°42'N
36°39'N
0 1 2 km
24°21'E
24°24'E
24°27'E
24°30'E
24°33'E

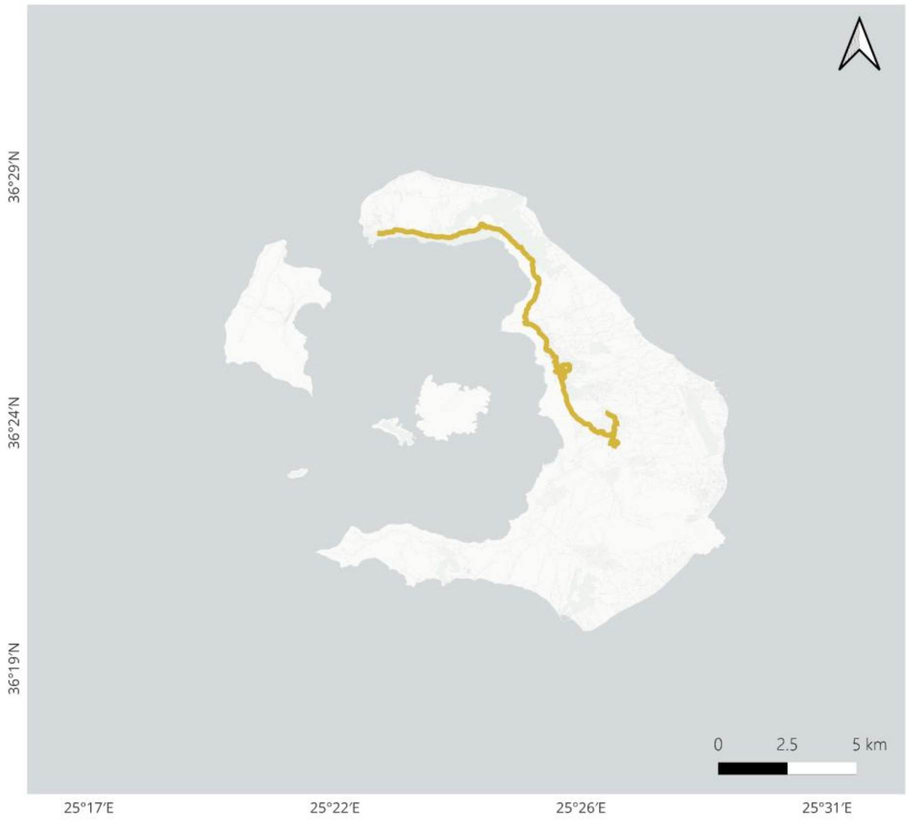
36°29′N
36°24′N
36°19′N
0
2.5
5 km
25°17′E
25°22′E
25°26′E
25°31′E

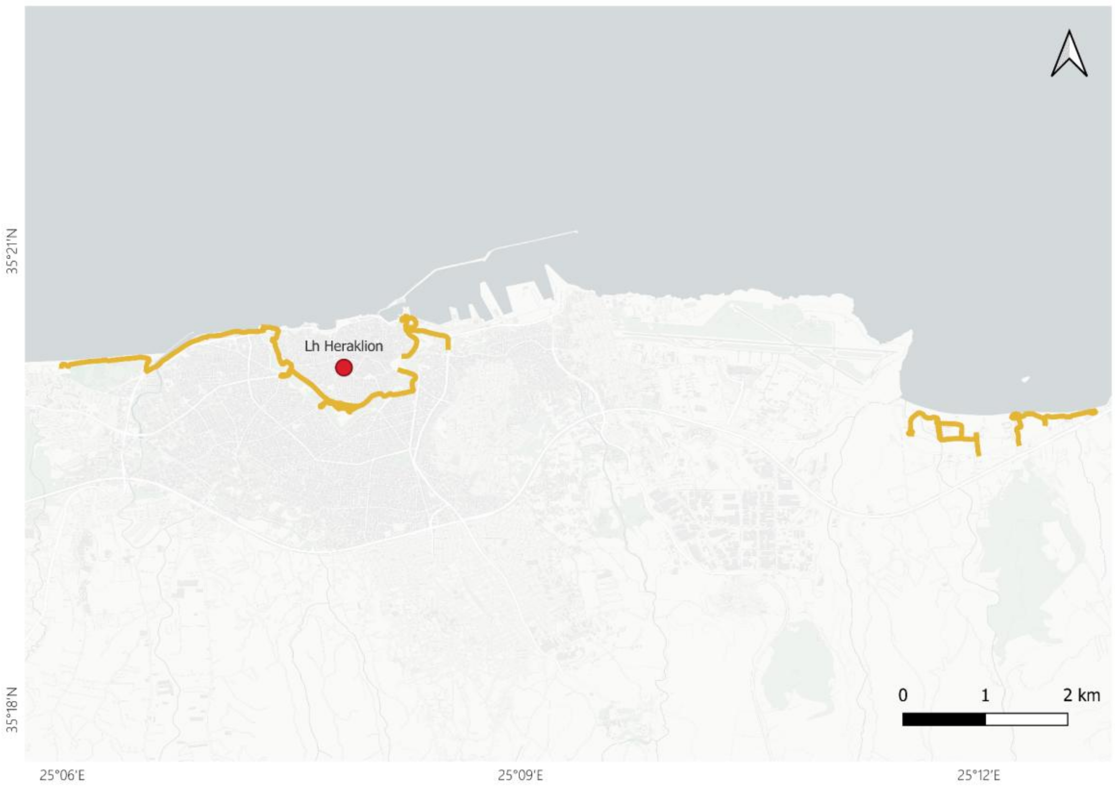
Lh Heraklion
35°21′N
35°18′N
0
1
2 km
25°06′E
25°09′E
25°12′E

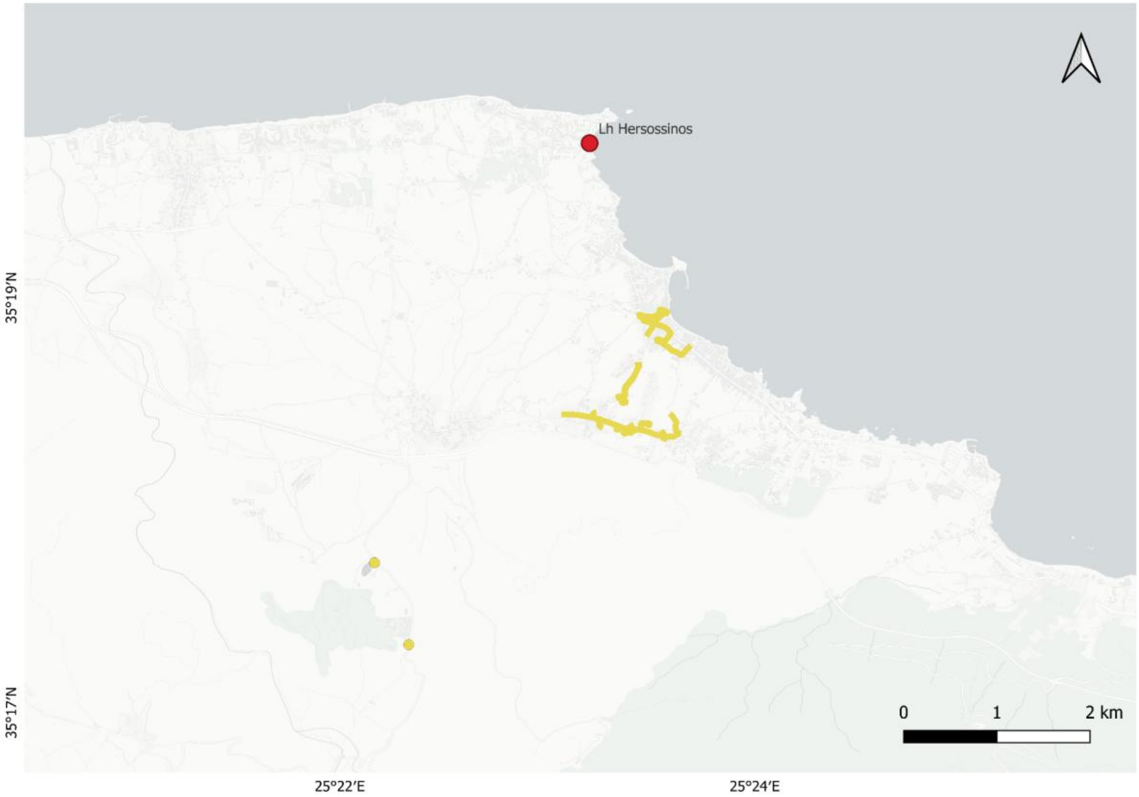


**Figure S3.** Barplots of Bayesian clustering results from the hierarchical approach based on the deltaK of Evanno, either without prior information (**A**) or with prior information based on colony membership (**B**).

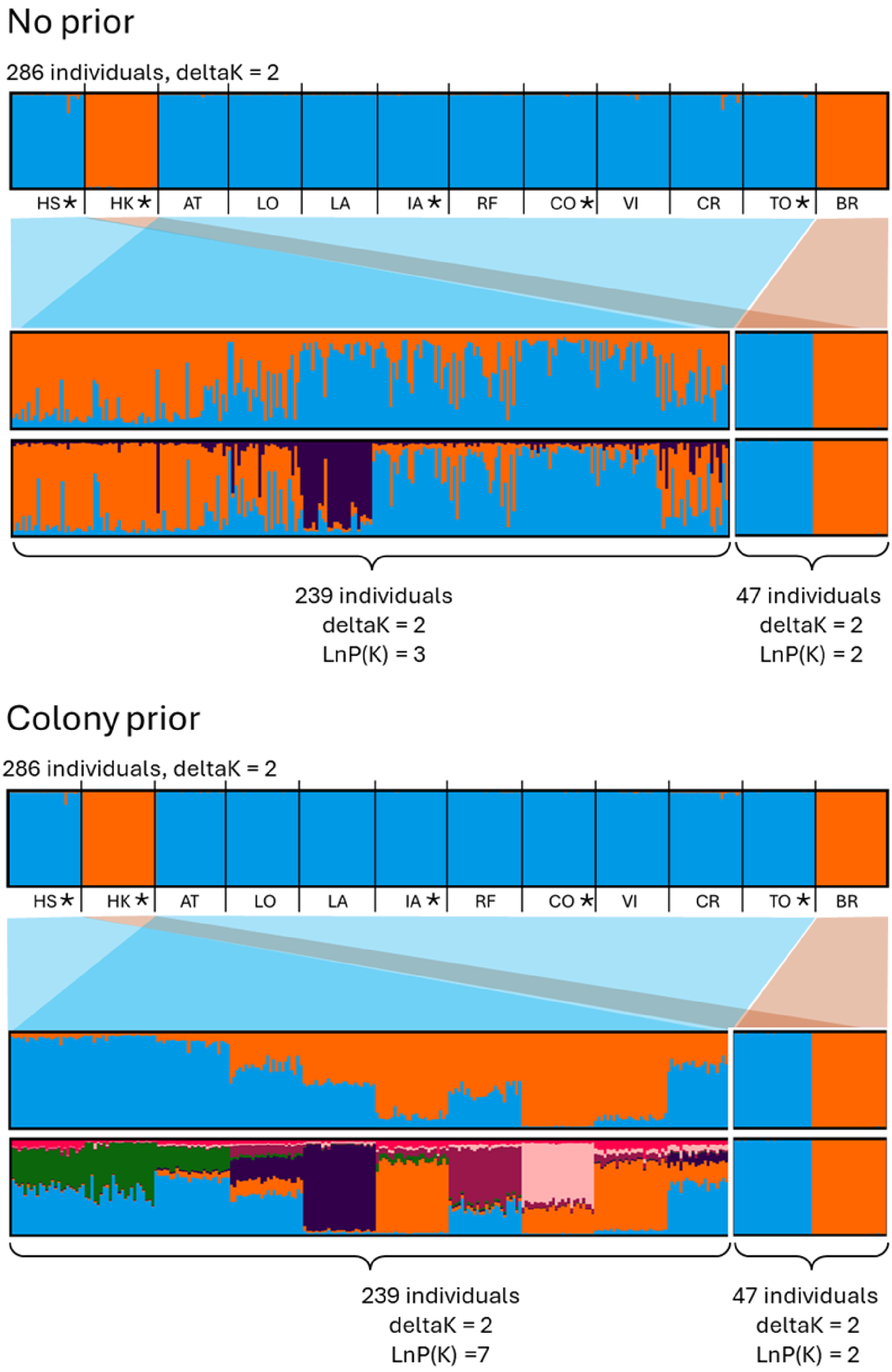
No prior
286 individuals, deltaK = 2
HS★
HK★
AT
LO
LA
IA★
RF
CO★
VI
CR
TO★
BR
239 individuals
deltaK = 2
LnP(K) = 3
47 individuals
deltaK = 2
LnP(K) = 2
Colony prior
286 individuals, deltaK = 2
HS★
HK★
AT
LO
LA
IA★
RF
CO★
VI
CR
TO★
BR
239 individuals
deltaK = 2
LnP(K) =7
47 individuals
deltaK = 2
LnP(K) = 2
